# Strand-independent degradation of uncoupled forks by EXO1 activates ATR and restrains fork progression

**DOI:** 10.64898/2026.03.22.666038

**Authors:** Emma J Grogan, Obehi E Ozua, Tamar Kavlashvili, Sara C Conwell, James M. Dewar

## Abstract

Nascent DNA degradation of replication forks shapes genome stability, yet the mechanism of degradation, and its direct consequences, remain unclear. Here we use localized nascent strand degradation at replication forks in *Xenopus* egg extracts to examine the mechanism and consequences of degradation. The exonuclease EXO1 is crucial for degradation and acts specifically on replication fork structures. Degradation requires EXO1 catalytic activity and is defective in the E109K Lynch Syndrome associated mutant. EXO1 degrades both nascent strands 5’-3’: the lagging strand from its native 5’ end and the leading strand from a distal 5’ entry on the sister fork, whereas leading strand 3’ end remains stable. Impaired leading-strand degradation at a specific site does not affect degradation of the corresponding lagging-strand region, so the two strands are degraded independently. Degradation of the uncoupled fork has two downstream consequences. It is important to activate ATR, which is otherwise weakly activated by the uncoupled fork. Additionally, degradation restrains fork progression, independent of ATR activation. Our findings demonstrate that strand-independent degradation of uncoupled forks by EXO1 activates ATR and restrains fork progression

## Introduction

Nascent DNA at stalled replication forks is vulnerable to nascent strand degradation and this is tightly regulated to preserve fork integrity^1–5^. Nascent strand degradation is implicated in genome instability^6–9^, innate immune signaling^10–12^, developmental abnormalities^13,14^, and progeria^15^. Excessive degradation is strongly correlated with sensitivity to DNA damaging agents^16–18^ and PARP inhibitors^19–27^, though this may be indirect^24,28^. Where a direct relationship between extensive resection and cancer cell viability is established, it can either cause cancer cell killing^29^ or be required for cancer cell viability^30^. Understanding the roles that nascent strand degradation plays in genome stability, cell viability, and human disease is therefore important. However, key questions remain about the mechanisms of degradation and the immediate downstream consequences.

Nascent strand degradation acts on replication forks^30–33^ and post-replicative ssDNA gaps^7,30–37^. Degradation is predominantly measured by fiber analysis^4^, where signal loss reflects loss of both the leading and lagging strands^38^, and would be expected to proceed 3’-5’ and 5’-3’ to degrade the leading and lagging strands, respectively. Yet the nucleases implicated in fork degradation act predominantly 5’-3’^30,31,34,35,39–41^, while 3’-5’ exonuclease activities implicated generally act to prevent degradation^6,12,34,42,43^. The exception is MRE11, which can perform 3’-5’ degradation at gaps^31,32^. However, it is unclear whether MRE11 performs 3’-5’ degradation at forks, given that its primary role is to initiate 5’-3’ degradation^3^, and the 3’ ends should be protected by RFC-PCNA^44,45^. One possibility is that the same mechanism operates on both leading and lagging strands, suggested by analysis of degradation at gaps^31^, but not directly demonstrated. Additionally, a range of protection factors induce degradation of both strands when inhibited^46^, suggesting that degradation of the two strands is interdependent, but this remains untested. Thus, it remains unclear how degradation of both strands occurs, the relative stability of the 5’ and 3’ ends, and to what extent degradation of the two strands is interdependent.

Both pathological and beneficial cellular roles are ascribed to nascent strand degradation. However, direct attribution is complicated by the fact that the same nucleases can operate at both forks^30–33^ and gaps^3,7,34–37^, while also participating in other genome stability pathways^3^. The strongest functional support is for a pathological role^7,24^ that causes genome instability. A beneficial role was reported for 3’-5’ exonuclease excision of chain-terminating or modified nucleotides ^47,48^ though whether this relates to extensive degradation is unclear. Roles in promoting^34,35,49–51^ and inhibiting^14^ replication restart and progression^52^ were reported but largely rely on the same nascent strand analysis used to detect degradation, leaving it unclear to what extent altered nascent strands reflect altered fork progression versus degradation. Degradation is also reported to promote activation of the ATR checkpoint^30,53^ yet, under the conditions where this occurs, helicase-polymerase uncoupling should already generate the DNA structures required for robust ATR activation^54^. Given that the ATR checkpoint regulates replication^55,56^, and may directly regulate fork progression^57,58^, it also raises the question of whether any effects of degradation on replication are direct or indirect effects of ATR signaling. Thus, direct outcomes of nascent strand degradation are unclear, making it difficult to understand its contribution to genome stability.

Here we elucidate a mechanism by which uncoupled forks are degraded and determine the direct consequences. Using *Xenopus* egg extracts, we examined synchronous, localized nascent strand degradation that allows us to directly monitor the degradation of individual DNA strands. We found that the 5’-3’ exonuclease EXO1 is critical for degrading both DNA strands of uncoupled forks but not reversed forks. Degradation requires the catalytic activity of EXO1 and is lost in the Lynch-syndrome E109K variant^59^. Lagging strands are degraded 5’-3’ starting at the 5’ end, while leading strands are degraded 5’-3’ from a distal 5’ entry point, which in this system is the lagging strand of the sister fork. We find that impaired leading-strand degradation does not affect lagging strand degradation, indicating that degradation of the two strands is independent. Surprisingly, we find that this degradation mechanism is crucial for ATR checkpoint activation even though the uncoupled fork itself should be sufficient. We further show that EXO1-dependent degradation restrains fork progression independent of its role in ATR activation. Overall, our data define a 5’-3’ degradation mechanism that works to degrade both strands independently in order to activate the ATR checkpoint and restrain fork progression, and is defective in the EXO1 Lynch-syndrome E109K variant^59^.

## Results

### EXO1 is crucial for nascent strand degradation

We previously characterized nascent strand degradation (NSD) at uncoupled replication forks that involved DNA2. We next tested the role of EXO1, the other long-range exonuclease involved in resection of replication forks^31,32,35,60–62^. We induced NSD as described^36^ in mock- or EXO1-immunodepleted *Xenopus* egg extracts; a LacR-bound *lacO* array localized synchronized forks to a defined region (Fig. 1A, Fig. S1A-E). Adding IPTG and the polymerase inhibitor aphidicolin generated structures corresponding to uncoupled replication forks (θ*; Fig. S1C-E), as described^36^. In mock-depleted extracts, this caused time-dependent loss of replication fork structures (RIs; Fig. 1B, lanes 1-5) and nascent strand signal (Fig. 1C), indicating robust degradation^36^. Immunodepletion of over 99% of EXO1 (Fig. S1A) stabilized fork structures and gave no detectable degradation (Fig. 1B, lanes 6-10; Fig. 1C). Monomeric daughter molecules from complete unwinding also formed (lin; Fig. 1B), indicating EXO1 normally restrains fork progression (Fig. 6, below). Degradation was rescued by purified recombinant EXO1 (Fig. 1B, lanes 11-15; Fig. S1B). Uncoupling kinetics were unaltered across all conditions, excluding impaired uncoupling as the cause of stability without EXO1 (Fig. S1E). Fork structures were slightly more compact in the rescue than mock at late time points (Fig. S1D, lanes 5 and 15), suggesting a minor recombinant-protein effect on topology, but were otherwise very similar. Thus, EXO1 is crucial for nascent strand degradation upon aphidicolin-induced uncoupling and also restrains fork progression.

**Figure 1:**
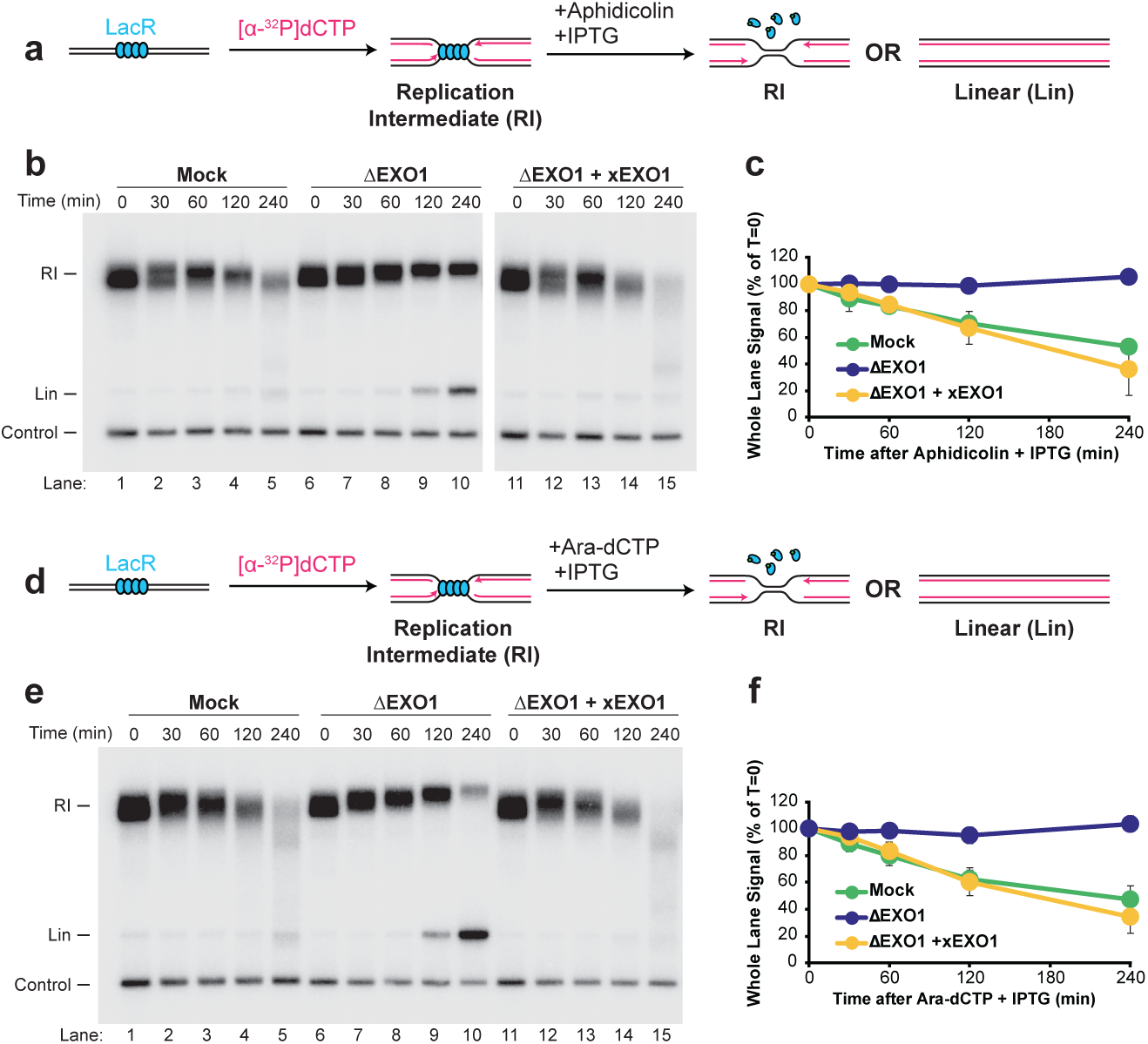
EXO1 is crucial for NSD in response to DNA polymerase stalling. **a,** Plasmid DNA harboring a LacR array was replicated in mock- and EXO1-immunodepleted *Xenopus* egg extracts in the absence or presence of purified recombinant xEXO1 protein, with [α-^32^P]dCTP added to label newly synthesized strands. Once forks were localized to the LacR barrier, aphidicolin and IPTG were added simultaneously to induce uncoupling. Samples were purified and digested with XmnI to allow unambiguous identification of replication intermediates (RI) and linear (Lin) products of replication. **b,** Samples from (a) were separated on an agarose gel and visualized by autoradiography. A plasmid lacking a LacR array was included as a loading control (Control). Representative of three replicates. **c,** Quantification of whole-lane signal from (b). Mean ± s.d., n = 3. **d,** As in (a), but with ara-dCTP in place of aphidicolin to stall nascent strands. **e,** Samples from (d) were separated on an agarose gel and visualized by autoradiography, with a plasmid lacking a LacR array as a loading control (Control). Representative of three replicates. **f,** Quantification of whole-lane signal from (e). Mean ± s.d., n = 3.

We asked whether the EXO1 requirement extended beyond aphidicolin to other means of uncoupling. We used ara-dCTP, a chain-terminating analog validated for stalling DNA polymerases in *Xenopus* egg extracts^63^, in the triphosphate form to avoid ara-C-conversion variability. We induced uncoupling and NSD as before but with ara-dCTP (Fig. 1D). As with aphidicolin, ara-dCTP induced robust degradation (Fig. 1E, lanes 1-5; Fig. 1F) and rapid uncoupling (Fig. S1F-H). Fork structures were restored later (θs; Fig. S1G, lane 5), indicating recovery from ara-dCTP stalling, unlike the permanent uncoupling with aphidicolin (Fig. S1D, lane 5). EXO1 immunodepletion diminished degradation and stabilized fork structures (RIs; Fig. 1E, lanes 6-10; Fig. 1F) without altering uncoupling kinetics (Fig. S1G, lanes 6-10), while permitting monomeric daughter molecules (lin), indicating EXO1 normally restrains fork progression, as for aphidicolin (Fig. 1A-C). Recombinant EXO1 reversed these effects (Fig. 1E, lanes 11-15, Fig. 1F). Again, the rescue migrated slightly faster than mock (Fig. S1G, lanes 5 and 15) but was otherwise similar. Thus, EXO1 is also required for NSD and restrains forks following ara-dCTP-induced uncoupling (Fig. 1D-F). With the aphidicolin data (Fig. 1A-C), EXO1 is crucial for NSD following fork uncoupling and restrains uncoupled-fork progression.

### EXO1 degrades uncoupled forks

We previously found that both uncoupled and reversed forks are degraded during NSD. To determine which structure EXO1 acts upon, we used 2D gel electrophoresis to analyze the structures formed in mock- or EXO1-immunodepleted extracts (Fig. 2A). In mock-treated extracts, uncoupled forks (double-Ys, DYs; Fig. 2B; Fig. S2A) initially declined while reversed forks (RFs; Fig. 2B; Fig. S2B) concurrently increased, as described^36^. The DYs then gained electrophoretic mobility, corresponding to degradation, and lost signal (Fig. 2Bii-iv, Fig. 2C), whereas RFs became more diffuse but did not appreciably decline (Fig. 2D). This differs from our previous results, where RFs also degraded, and reflects a nonspecific effect of immunodepletion, either creating asynchrony in RF formation and degradation or inactivating degradation. We nonetheless observed extensive NSD and DY-to-RF conversion, as described^36^.

**Figure 2:**
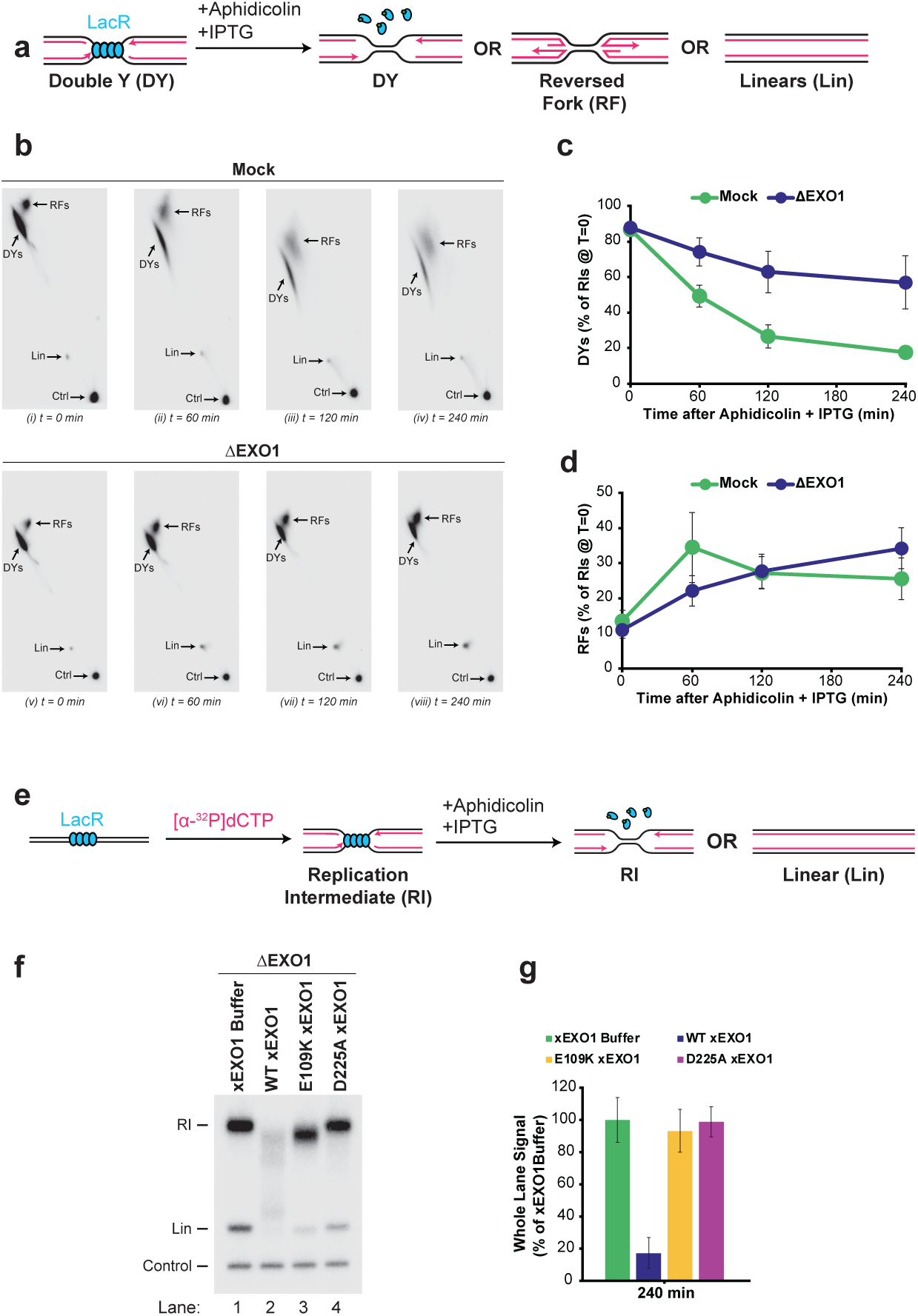
EXO1 degrades uncoupled forks, not reversed forks. **a,** Plasmid DNA harboring a LacR array was replicated in mock- and EXO1-immunodepleted *Xenopus* egg extracts. Once forks were localized to the LacR barrier, aphidicolin and IPTG were added simultaneously to induce uncoupling. Samples were purified and digested with XmnI to yield double-Ys (DYs), reversed forks (RFs), and linear products of replication (Lin). **b,** Samples from (a) were separated by 2D gel electrophoresis and visualized by autoradiography, with a plasmid lacking a LacR array as a loading control (Control). Representative of three replicates. **c,** Quantification of DYs from (b). Mean ± s.d., n = 3. **d,** Quantification of RFs from (b). Mean ± s.d., n = 3. **e,** Plasmid DNA harboring a LacR array was replicated in EXO1-immunodepleted *Xenopus* egg extracts supplemented with EXO1 wild-type, E109K, or D225A protein, with [α-^32^P]dCTP added to label newly synthesized strands. Once forks were localized to the LacR barrier, aphidicolin and IPTG were added simultaneously to induce uncoupling, and samples were purified and digested as in Fig. 1a. **f,** Samples from (e) were separated on an agarose gel and visualized by autoradiography. Representative of three replicates. **g,** Quantification of whole-lane signal from (f) relative to the buffer condition. Mean ± s.d., n = 3.

We next examined structures formed when EXO1 was immunodepleted (Fig. 2Bv-viii). Uncoupled forks still decreased initially (DYs; Fig. S2A) and reversed forks increased (RFs; Fig. S2B), but less extensively than in mock-treated extracts. We also unambiguously detected fully unwound linear molecules (Lin; Fig. 2Bv-vi, Fig. S2C), supporting increased fork progression upon EXO1 removal. Most uncoupled-fork signal persisted (Fig. 2C) with little change in mobility (DYs; Fig. 2Bv-vi), indicating little degradation. Reversed-fork abundance also differed little between mock- and EXO1-depleted extracts (RFs; Fig. 2Bii,vi, Fig. 2D). Thus, degradation by EXO1 occurred predominantly at uncoupled forks. Because reversed-fork mobility was not detectably altered (Fig. 2Bv-viii), we cannot discern whether this reflects no direct role for EXO1 in degrading reversed forks or an indirect consequence of reduced degradation of their precursors. Overall, EXO1 is important for degradation of uncoupled forks.

Previously reported forms of EXO1-mediated NSD are positively or negatively regulated by the homologous recombination protein RAD51^7,35,37^, so we asked whether RAD51 was involved in degradation of uncoupled forks (Fig. S2D). Immunodepletion of RAD51 (Fig. S2E) produced no detectable change in the extent of NSD (Fig. S2F-G). Thus, EXO1-dependent degradation of uncoupled forks is distinct from the RAD51-regulated degradation pathways.

### Enzymatic and non-enzymatic roles of EXO1

EXO1 has a well-defined exonuclease activity inactivated by the D225A mutation in the catalytic core^64^ and a proposed non-enzymatic function inactivated by the E109K Lynch syndrome mutation^59,65,66^. To test which functions degradation requires, we purified *Xenopus* EXO1 carrying each mutation (Fig. S2H) and compared it to wild type in EXO1-immunodepleted extracts. Neither mutation appreciably affected replication efficiency (Fig. S2I). As expected, wild type restored degradation of replication fork structures (RIs; Fig. 2F, lane 2, Fig. S2J), but neither the D225A nor the E109K mutant did (Fig. 2F, lanes 3-4). Our usual normalization to signal at degradation onset was highly variable (Fig. S2J); normalizing only to our internal control (Fig. 2F) instead showed clear, consistent degradation by wild type but not either mutant (Fig. 2G), with signal consistent across conditions before degradation (Fig. S2K). We conclude that degradation requires both the catalytic activity of EXO1 and the non-enzymatic function defective in the Lynch syndrome E109K mutant.

### 5’-3’ degradation of leading and lagging strands

To determine the polarity of leading- and lagging-strand degradation, we generated a plasmid with nicking sites positioned proximal and distal to the 3’ and 5’ ends of the nascent leading and lagging strands (strand analysis plasmid; Fig. S3A). The leading- and lagging-strand sites were at the same location but on opposite strands, allowing direct comparison within the same region of the nascent duplex (Fig. S3A). Digestion also released a fragment from a region fully replicated before NSD induction, serving as an internal loading control (control plasmid; Fig. S3A). This allowed us to monitor degradation proximal to the leading 3’ end or the lagging 5’ end versus distal to either end.

We first examined nascent lagging strands after NSD induction (Fig. 3A). Signal proximal to the 5’ end was rapidly lost (lag end; Fig. 3B-C, Fig. S3B-C); most disappeared within the first 60 minutes and was essentially undetectable by 120 minutes. Degradation distal to the 5’ end was delayed (lag internal; Fig. 3B-C), occurring mostly between 60 and 120 minutes and becoming undetectable by 240 minutes. Faster loss nearer the 5’ end demonstrates 5’-3’ degradation, consistent with the crucial role for the 5’-3’ exonuclease EXO1 (Fig. 1).

**Figure 3:**
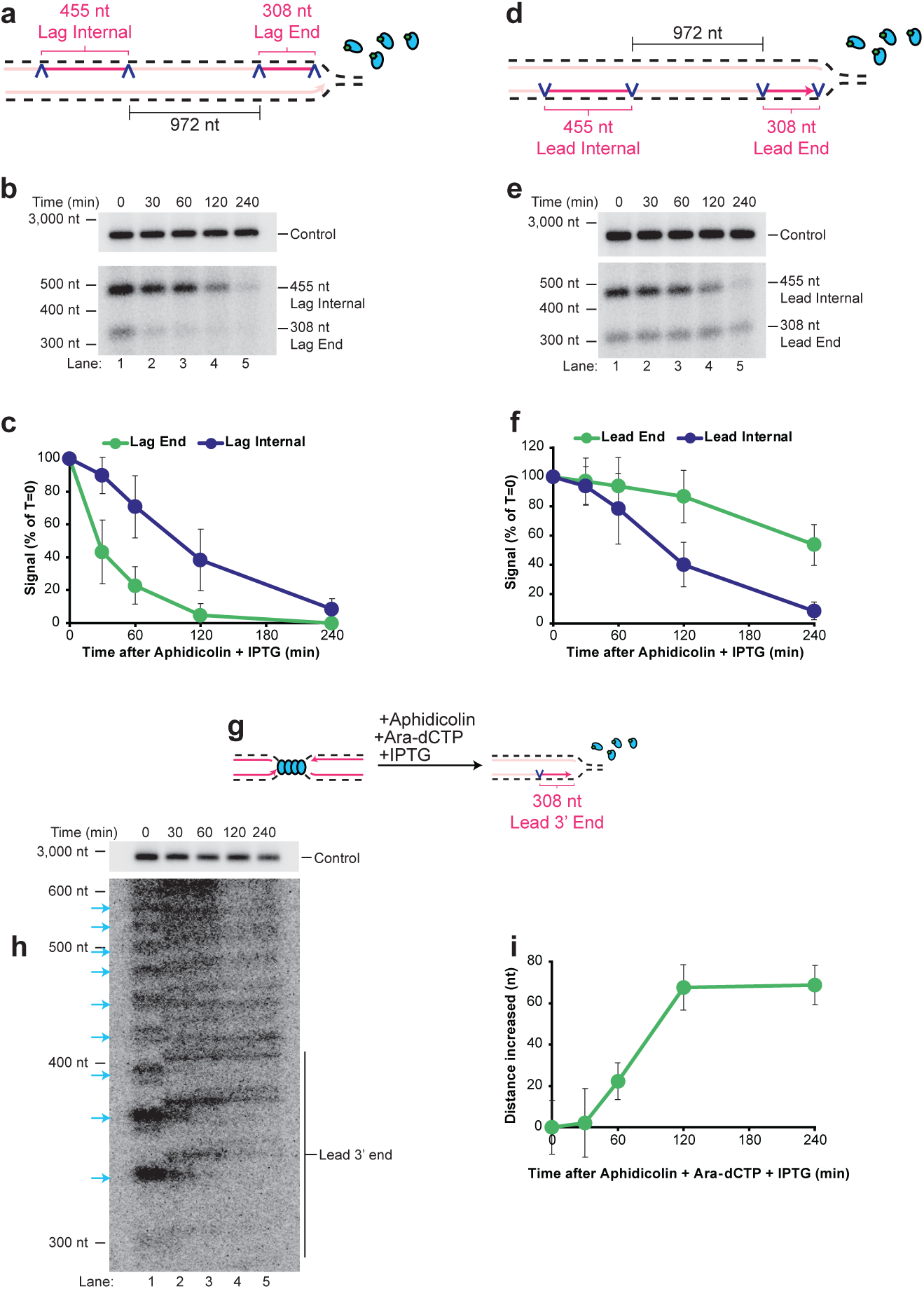
Both leading and lagging strands are degraded 5’-3’. **a,** Plasmid DNA harboring a LacR array was replicated and uncoupled in *Xenopus* egg extracts as in Fig. 1a. Samples were purified and digested with SacI and Nb.BbvCI to release two nascent lagging-strand fragments of different size and position: a 308 nt fragment at the 5’ end of the nascent lagging strand and a 455 nt fragment internal to the nascent lagging strand. **b,** Samples from (a) were separated on native (top) and denaturing (bottom) agarose gels and visualized by autoradiography. Representative of three replicates. **c,** Quantification of lagging-strand end and internal fragments from (b). Mean ± s.d., n = 3. **d,** As in (a), but digested with SacI and Nt.BbvCI to release two nascent leading-strand fragments: a 308 nt fragment at the 3’ end of the nascent leading strand and a 455 nt fragment internal to the nascent leading strand. **e,** Samples from (d) were separated on native (top) and denaturing (bottom) agarose gels and visualized by autoradiography. Representative of three replicates. **f,** Quantification of leading-strand end and internal fragments from (e). Mean ± s.d., n = 3. **g,** Plasmid DNA harboring a LacR array was replicated in *Xenopus* egg extracts. Once forks were localized to the LacR barrier, aphidicolin, ara-dCTP, and IPTG were added simultaneously to induce uncoupling. Samples were purified and digested with Nt.BbvCI to monitor the nascent leading-strand 3’ end. **h,** Samples from (g) were separated on native (top) and denaturing polyacrylamide (bottom) gels. Arrows indicate stalling at distinct LacR-bound *lacO* sites (as in ^69^). Representative of three replicates. **i,** Quantification of the increase in leading-strand 3’ end distance over time from (h). Mean ± s.d., n = 3.

We next examined nascent leading strands (Fig. 3D). Surprisingly, signal proximal to the 3’ end was relatively stable (lead end; Fig. 3E-F, Fig. S3D-E), with no detectable loss at 120 minutes and approximately 50% remaining at 240 minutes. Degradation distal to the 3’ end was faster (lead internal; Fig. 3E-F), following similar kinetics to the lagging strand distal to the 5’ end (Fig. 3B-C). This similarity is expected, because the distal regions lie opposite each other at comparable distances from the two forks (1280 bp and 1144 bp; Fig. S3A), each roughly equidistant from a leading-strand 3’ end and a lagging-strand 5’ end. Greater degradation of internal sequences than the 3’ end indicates leading-strand degradation also proceeds 5’-3’, and the prolonged 3’-end stability indicates these ends undergo limited degradation.

To more sensitively analyze stability of 3’ ends, we analyzed their abundance and position by denaturing alkaline gel electrophoresis (Fig. S3L) and found they were slowly extended even with aphidicolin (Fig. S3M-N) at ∼25 nts per hour, which could mask degradation. We therefore combined aphidicolin and ara-dCTP, which both induce EXO1-dependent NSD (Fig. 1) through distinct mechanisms^67,68^, and used higher-resolution denaturing polyacrylamide gel analysis to detect smaller nascent strands (Fig. 3G). Before NSD induction, we detected discrete products corresponding to forks stalled at different LacR-bound *lacO* sites (Fig. 3H, arrows), as described^69^. After induction, these products slowly increased in size (Fig. 3H), and we did not detect shorter products that would be expected from low-level degradation (Fig. S3F-K). Thus, we detected no products of 3’-5’ degradation.

3’-5’ degradation might still occur but be masked by rapid 5’-3’ resynthesis. To test this, we performed a pulse-chase, adding excess unlabeled deoxynucleotides at the onset of degradation (Fig. S3O), so degraded signal could not be restored by new synthesis. Leading-strand stability was largely unaffected, the 3’ end signal remaining stable for at least two hours (Fig. S3P-R). The chase blocked over 95% of radiolabel incorporation (Fig. S3S-U), and synthesis quantification showed net extension of approximately 70 nucleotides over four hours (Fig. 3I), or ∼17.5 nts per hour. Thus, 3’ ends are largely stable and undergo slow net extension, even with two different DNA polymerase inhibitors.

### Strand-independent degradation by EXO1

Given that EXO1 was responsible for most nascent strand degradation (Fig. 1), we examined its specific contribution to the 5’-3’ degradation on both the leading and lagging strands.

We first immunodepleted EXO1 and examined lagging strand degradation (Fig. 4A). In mock-immunodepleted extracts, signal proximal to the 5’ end was rapidly lost, while distal signal was lost more slowly and to a lesser extent (Fig. 4B-D, Fig. S4A-B), as observed previously (Fig. 3B-C). Here the distal signal was more stable than in earlier experiments, likely due to a nonspecific reduction in extract activity from the depletion procedure (compare Fig. 4D mock with Fig. 3C). Strikingly, EXO1 immunodepletion ameliorated most signal loss, both proximal and distal to the 5’ end (Fig. 4B-D). Although some degradation occurred, there was no detectable difference between proximal and distal signal (Fig. S4D), showing the 5’-3’ degradation was lost. EXO1 is therefore crucial for 5’-3’ lagging strand degradation.

**Figure 4:**
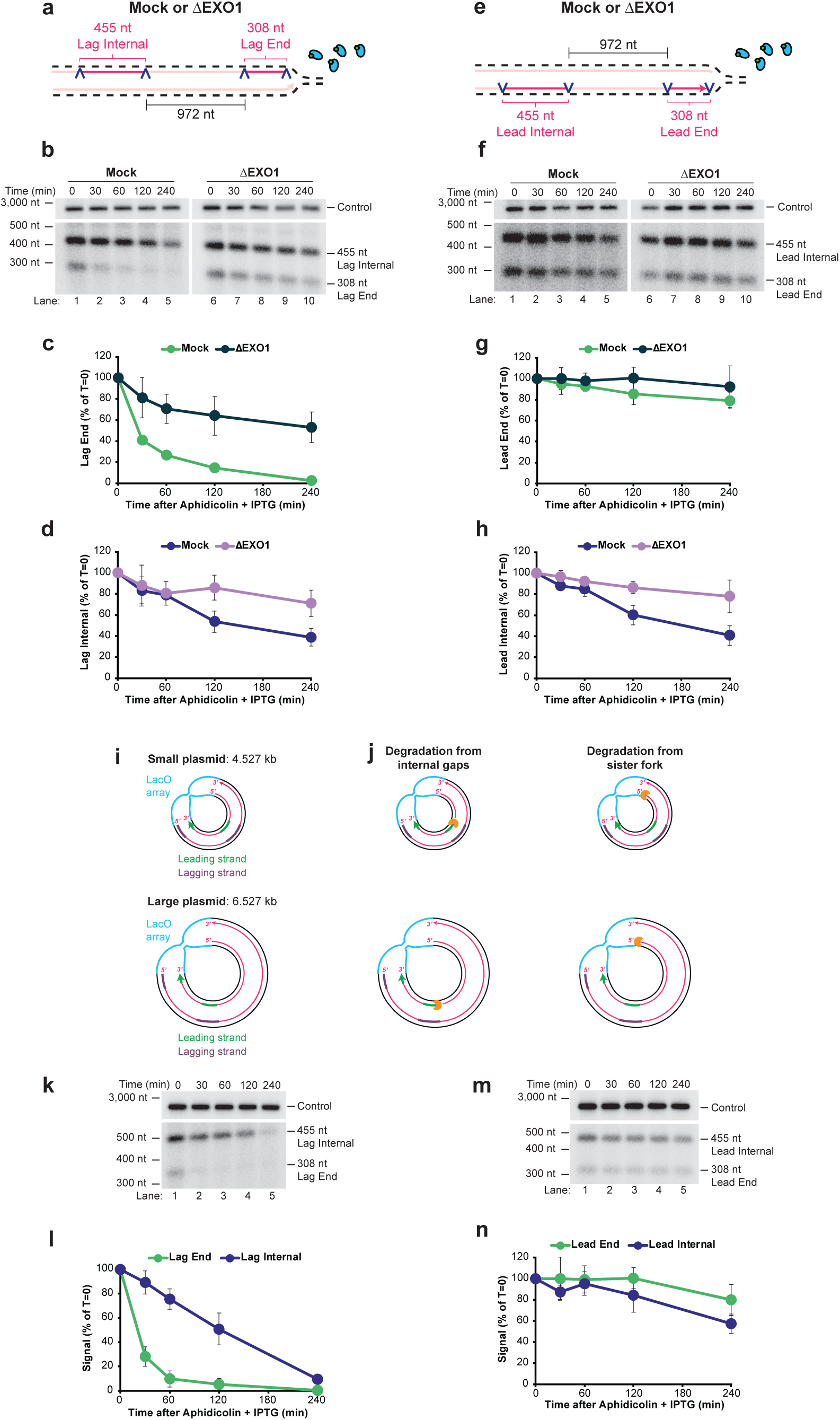
EXO1 mediates strand-independent degradation. **a,** Plasmid DNA harboring a LacR array was replicated and uncoupled in mock- and EXO1-immunodepleted *Xenopus* egg extracts as in Fig. 3a, and digested as in Fig. 3a. **b,** Samples from (a) were separated on native (top) and denaturing (bottom) agarose gels and visualized by autoradiography. Representative of three replicates. **c,** Quantification of the lagging-strand end fragment from (b). Mean ± s.d., n = 3. **d,** Quantification of the lagging-strand internal fragment from (b). Mean ± s.d., n = 3. **e,** Plasmid DNA harboring a LacR array was replicated and uncoupled in mock- and EXO1-immunodepleted *Xenopus* egg extracts as in Fig. 3d, and digested as in Fig. 3d. **f,** Samples from (e) were separated on native (top) and denaturing (bottom) agarose gels and visualized by autoradiography. Representative of three replicates. **g,** Quantification of the leading-strand end fragment from (f). Mean ± s.d., n = 3. **h,** Quantification of the leading-strand internal fragment from (f). Mean ± s.d., n = 3. **i,** Cartoon of the two plasmid templates used to monitor strand-specific degradation. The small template used in Fig. 3a-f is 4.527 kb; the large template is 6.527 kb and carries an additional 2 kb of sequence between the internal fragments and the converging sister fork. **j,** Cartoons of the predicted outcomes for the internal-gaps hypothesis (left) and the sister-fork hypothesis (right) for leading-strand internal degradation. **k,** Digestion was performed as in Fig. 3a but using the large template; samples were separated on native (top) and denaturing (bottom) agarose gels and visualized by autoradiography. Representative of three replicates. **l,** Quantification of lagging-strand end and internal fragments from (k). Mean ± s.d., n = 3. **m,** Digestion was performed as in Fig. 3d but using the large template; samples were separated on native (top) and denaturing (bottom) agarose gels and visualized by autoradiography. Representative of three replicates. **n,** Quantification of leading-strand end and internal fragments from (m). Mean ± s.d., n = 3.

We then examined leading strand degradation (Fig. 4E). In mock-immunodepleted extracts, signal proximal to the 3’ end was more stable than distal signal (Fig. 4F-H, Fig. S4E-F), consistent with prior observations (Fig. 3E-F). The leading strands were more stable than before, again likely reflecting reduced extract activity from the depletion (compare Fig. 4G-H mock with Fig. 3F). EXO1 immunodepletion had a negligible effect proximal to the 3’ end (Fig. 4G), consistent with the little degradation of 3’ ends (Fig. 3) but ameliorated most degradation distal to the 3’ end (Fig. 4H). As for lagging strands, loss of EXO1 eliminated the difference between proximal and distal sequences (Fig. S4H), showing 5’-3’ degradation was lost. EXO1 was thus also crucial for 5’-3’ leading strand degradation.

EXO1’s requirement for both strands raised the question of leading strand degradation. Given its 5’-3’ polarity, degradation could initiate either at internal gaps on the leading strand^31,32^ or at the 5’ end of the nascent lagging strand on the sister fork. To distinguish these, we generated a plasmid with a larger backbone, increasing the distance between the converging forks’ start sites from ∼3,000 bp to ∼5,000 bp (Fig. 4I). Internal-gap degradation should be largely unaffected by this distance, whereas sister-fork-initiated degradation should be substantially diminished by the greater travel distance (Fig. 4J). Lagging strand degradation was largely unaffected by sister-fork position (Fig. 4K-L, Fig. S4I-L) as expected for 5’-3’ degradation initiating at the 5’ end. In contrast, nascent leading strands were substantially stabilized, with most degradation now blocked (Fig. 4M-N, Fig. S4M-P), consistent with most leading strand degradation initiating from the sister fork. This data also demonstrates independence of strand degradation: because the increased distance diminished leading-strand degradation without affecting the lagging strand, lagging-strand degradation does not require leading-strand degradation. Overall, EXO1 degrades leading and lagging strands independently starting from a 5’ end and proceeding 5’-3’.

### EXO1 activates ATR at uncoupled forks

EXO1 activity reportedly promotes ATR checkpoint activation at DNA gaps^30^, so we tested whether EXO1 activity at uncoupled forks also promotes ATR activation. Our NSD approach (Fig. 1A) involves normal replication, fork stalling, and uncoupling, each of which could influence ATR signaling^70–72^. We therefore isolated these events and analyzed phosphorylation of the ATR substrate CHK1 as a readout for ATR activation.

We first tested normal replication and fork stalling. Plasmid DNA was replicated without a LacR barrier or with LacR to stall forks (Fig. 5A). Without LacR, replication fork structures were detectable by 5 minutes and declined by 20 minutes as replication completed (θs; Fig. 5B, lanes 1-2), with circular monomeric plasmids predominating at 20 minutes (Fig. 5B, lane 2). With LacR, forks stalled stably, leaving only fork structures from 20 to 80 minutes (θs; Fig. 5B, lanes 3-4). Neither condition produced appreciable CHK1 phosphorylation (Fig. 5C, lanes 1-4).

**Figure 5:**
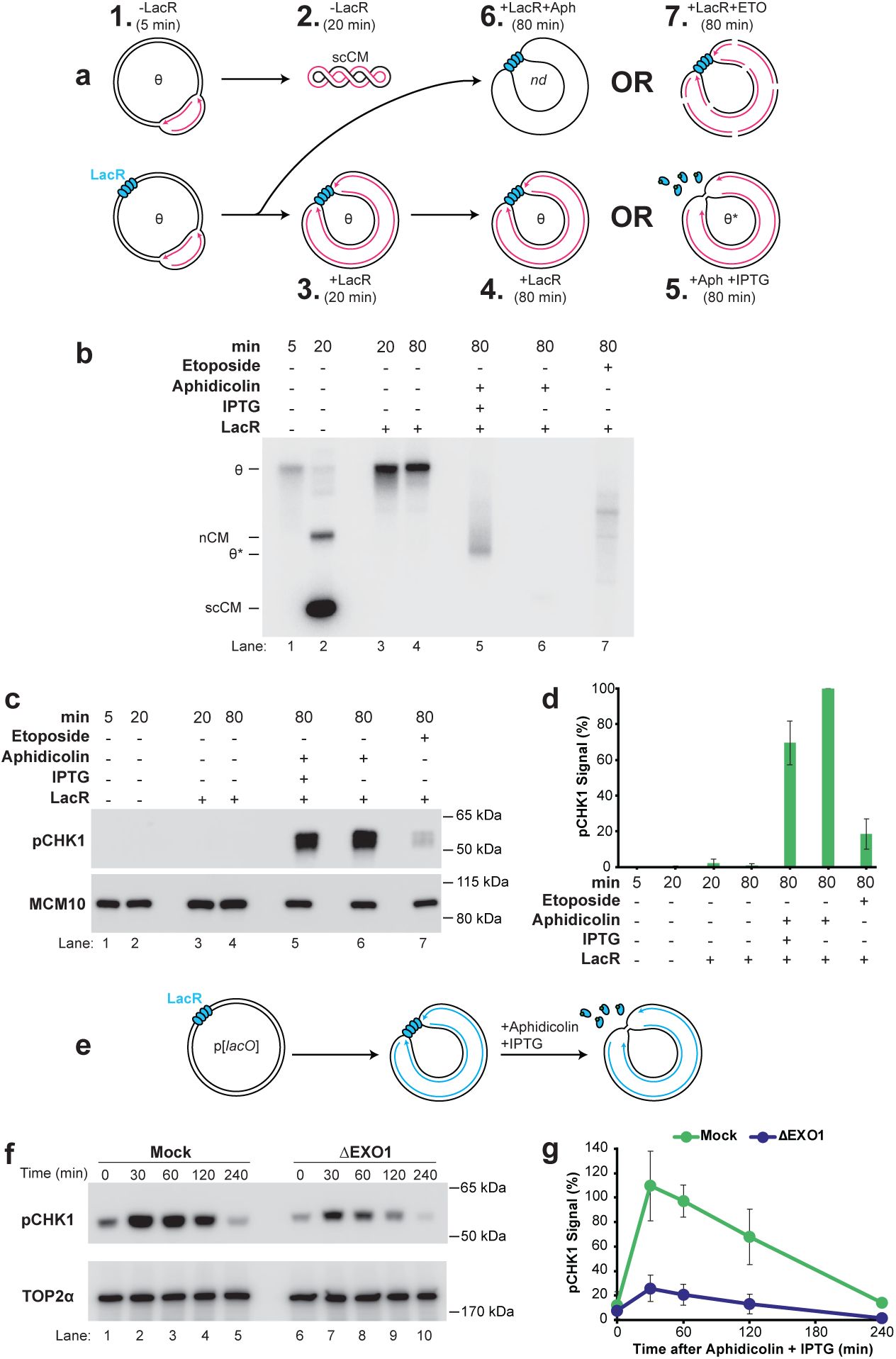
EXO1 degradation at uncoupled forks activates ATR. **a,** Plasmid DNA was replicated in *Xenopus* egg extracts with [α-^32^P]dATP to yield: (1) θ structures; and (2) supercoiled monomeric (scCM) products. In parallel, plasmid DNA harboring a LacR array was replicated with [α-^32^P]dATP to yield either: (3) θ structures, when forks were localized to the LacR barrier; (4) θ structures, from prolonged stalling at the LacR barrier; (5) θ* structures, after simultaneous addition of aphidicolin and IPTG to induce uncoupling; (6) extensively unwound template when aphidicolin was added at the onset of replication; and (7) DNA double-strand breaks, when replicated in the presence of etoposide (as in ^81^). Numbers correspond to the lanes in (b) and (c). **b,** Samples from (a) were separated on an agarose gel and visualized by autoradiography. Representative of three replicates. **c,** Replication was performed as in (a) but without [α-^32^P]dATP (see Supplemental Fig. 5a); pCHK1 and MCM10 were detected by western blotting. MCM10 serves as a loading control. **d,** Quantification of pCHK1 signal from (c). Mean ± s.d., n = 3. **e,** Plasmid DNA was replicated in mock- or EXO1-immunodepleted *Xenopus* egg extracts; forks were localized to the LacR barrier, and uncoupling was induced by simultaneous addition of aphidicolin and IPTG. **f,** pCHK1 and TOP2α from (e) were detected by western blotting. TOP2α serves as a loading control. Representative of three replicates. **g,** Quantification of pCHK1 signal from (f). Mean ± s.d., n = 3.

**Figure 6:**
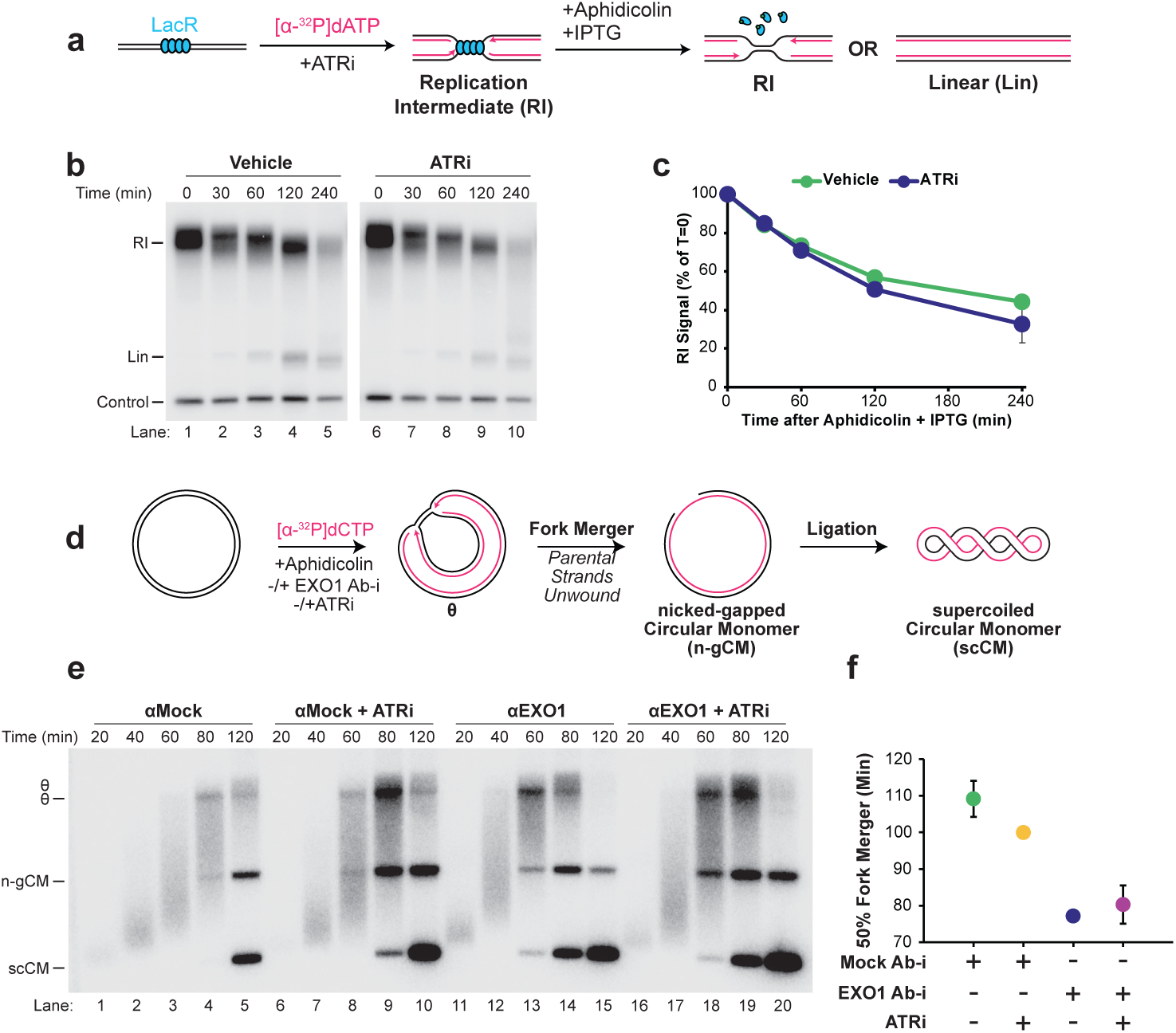
EXO1-mediated degradation of uncoupled forks restrains fork progression. **a,** Plasmid DNA harboring a LacR array was replicated in *Xenopus* egg extracts in the absence or presence of ATRi, with [α-^32^P]dATP added to label newly synthesized strands. Once forks were localized to the LacR barrier, aphidicolin and IPTG were added simultaneously to induce uncoupling. Samples were purified and digested as in Fig. 1a. **b,** Samples from (a) were separated on an agarose gel and visualized by autoradiography (RI, Lin, Control; vehicle, lanes 1-5; ATRi, lanes 6-10). Representative of three replicates. **c,** Quantification of RI signal from (b). Mean ± s.d., n = 3. **d,** Plasmid DNA was replicated in *Xenopus* egg extracts in the presence of aphidicolin, with mock or EXO1 antibody, and in the absence or presence of ATRi, with [α-^32^P]dCTP added to label newly synthesized strands. **e,** Samples from (d) were separated on an agarose gel and visualized by autoradiography, resolving θ, nicked, and supercoiled (SC) species at 20, 40, 60, 80, and 120 min (αMock, lanes 1-5; αMock + ATRi, lanes 6-10; αEXO1, lanes 11-15; αEXO1 + ATRi, lanes 16-20). Representative of three replicates. **f,** Quantification from (e) of the time in minutes at which fork merger was complete on 50% of molecules for each condition. Mean ± s.d., n = 3.

Uncoupling induced by aphidicolin addition at replication onset^71^ produced no detectable DNA synthesis (Fig. 5B, lane 6) but high CHK1 phosphorylation (Fig. 5C, lane 6). Our standard NSD protocol, localizing forks to the LacR barrier before simultaneous barrier release and aphidicolin addition (Fig. 5A), produced extensive uncoupling (Fig. 5B, lane 5) and high CHK1 phosphorylation (Fig. 5C, lane 5), within ∼30% of the levels from onset uncoupling (Fig. 5D). Thus, uncoupling potently stimulates CHK1 phosphorylation, as previously characterized^71^, but neither stalling nor unperturbed replication led to appreciable CHK1 phosphorylation.

To determine the specificity of checkpoint activation, we assessed ATM activation using ATM phosphorylation as a read-out (Fig. S5A-C). As a positive control we performed etoposide treatment, which generated the expected DNA fragmentation arising from DSBs (Fig. 5B, lane 7) and resulted in robust ATM activation (Fig. S5B lane 7, Fig. S5C) but only modest CHK1 phosphorylation (Fig. 5C, lane 7), as expected because DSBs stimulate ATM more than ATR. ATM activation was minimally affected by stalling or uncoupling relative to etoposide (Fig. S5A-C). Thus, the checkpoint response we observed was largely ATR-specific.

Having established that uncoupling as part of our NSD protocol activates CHK1, we tested the contribution of EXO1. We induced nascent strand degradation in mock- and EXO1-immunodepleted extracts and analyzed CHK1 activation over time (Fig. 5E). In mock-depleted extracts, CHK1 activation peaked shortly after uncoupling and then declined (Fig. 5F, lanes 1-5; Fig. 5G). Strikingly, EXO1 immunodepletion blocked most CHK1 activation (Fig. 5F, lanes 6-10; Fig. 5G). This was not an indirect effect of defective uncoupling, since EXO1 depletion did not alter uncoupling kinetics (Fig. S1D-E), nor a generalized requirement for EXO1 in ATR activation, because CHK1 phosphorylation was unaltered by EXO1 depletion when a model ATR-activating substrate was supplied (Fig. S5D-H). Thus, EXO1 activity at uncoupled forks is crucial for ATR activation.

### Overlapping but distinct roles for EXO1 and ATR

ATR can negatively regulate processing of uncoupled forks, so EXO1’s role in degradation could be partly an indirect effect of its role in ATR activation. To test this, we monitored nascent strand degradation with or without ATR inhibitor (ATR-i; used as in Fig. S5E). ATR-i had no appreciable effect on degradation of replication fork structures versus vehicle control (Fig. 6A-C). It also did not impact the generation of full length linear molecules (Fig. 6B), indicating that the fork restraint role of EXO1 was also not due to ATR. Thus, EXO1’s effect on degradation and restraint of uncoupled forks cannot be attributed to its role in ATR activation.

Our data that EXO1 restrains fork progression were generated under conditions where DNA synthesis was near-completely blocked and most forks did not fully unwind the DNA template (Fig. 1B,E, Fig. 2B, Fig. S2C). We therefore examined conditions where DNA synthesis was allowed to proceed in the presence of low, continual uncoupling induced by low-dose aphidicolin. Under these conditions, most fork structures resolved within an ∼40 minute window (θ; Fig. 6E, lanes 1-5; Fig. 6F), giving rise to the fully replicated supercoiled daughter molecules that are the final products of replication (SC; Fig. 6E, lanes 4-5). To test EXO1’s effect on uncoupled forks, we used an inhibitory antibody (EXO1-i) that blocked degradation by specifically inhibiting EXO1 (Fig. S6A-C); we chose this over immunodepletion to avoid asynchrony from nonspecific reductions in extract activity. EXO1-i accelerated disappearance of θs (Fig. 6E, lanes 11-15, Fig. 6F) and accumulation of SC (Fig. 6E, lanes 11-15; Fig. 6F), showing forks completed synthesis more quickly. This was not due to altered origin firing, as it persisted in a pulse-chase preventing detection of newly fired origins (Fig. S6E-H). The measurements were not confounded by altered nascent strand degradation because we measured resolution of replication structures (Fig. 6D-F), which is determined by unwinding of parental strands rather than nascent strand growth. Thus, EXO1 restrains fork movement following low levels of uncoupling, consistent with our NSD conditions (Fig. 1B,E, Fig. 2B, Fig. S2C).

The budding yeast analog of ATR signaling restrains replication forks^57^. We did not observe this under conditions where high levels of uncoupling were induced (linears were not produced in Fig. 6A-C) but it was possible that ATR contributed to the restraint of forks by EXO1 when lower levels of uncoupling were induced (Fig. 6D-F). To test this, we replicated DNA in the presence of low dose aphidicolin and added ATR inhibitor (ATRi; Fig. 6D). ATRi increased total replication signal (Fig. 6E, Fig. S6D), consistent with its role suppressing origin firing^72^. EXO1-i also increased total signal, non-additively with ATRi (Fig. 6E, Fig. S6D), indicating EXO1 contributes to ATR-mediated suppression of origin firing. ATRi also slightly hastened resolution of θs (Fig. 6E, lanes 6-10; Fig. 6F), suggesting ATR, like EXO1, restrained fork progression (Fig. 6F). However, this effect was less pronounced than EXO1 inhibition (Fig. 6F), and combined EXO1 and ATR inhibition matched EXO1 inhibition alone (Fig. 6E, lanes 16-20; Fig. 6F). Thus EXO1’s effect on fork progression may be partially dependent on ATR, whereas any effect of ATR on fork progression is entirely dependent on EXO1. This contrasts to the effect on total DNA synthesis, where EXO1’s effect is dependent on ATR, likely reflecting EXO1-mediated ATR activation. Overall, EXO1 restrains replication forks independently of ATR (Fig. 6F) while regulating total DNA synthesis in an ATR-dependent manner (Fig. S6D).

## Discussion

### Mechanism and consequences of nascent strand degradation by EXO1

Here we show that, following helicase-polymerase uncoupling, the 5’-3’ exonuclease EXO1 can degrade nascent DNA through its catalytic activity (Fig. 7A-C), consistent with its previously described role in degradation^31,32,35,60–62^. This degradation mechanism occurs at uncoupled forks rather than at the reversed forks into which they convert (Fig. 7E-F), and it proceeds independently of RAD51 (Fig. S2D-G). The RAD51-independence distinguishes processing of uncoupled forks from the canonical, RAD51-dependent degradation of reversed forks^7,35,37^. Both nascent strands are degraded with 5’-3’ polarity (Fig. 3), with initiation at 5’ ends (Fig. 4): lagging strands are degraded from their native 5’ ends, and leading strands from the 5’ end of the lagging strand of the sister fork. Degradation of the two strands is therefore strand-independent rather than coupled. This processing has two downstream consequences: it is crucial to activate the ATR-dependent checkpoint (Fig. 5, Fig. 7D); and it restrains fork progression (Fig. 6, Fig. 7D). Together with our previous finding that forks can be degraded without reversal^36^, we conclude that EXO1-dependent degradation and fork reversal are parallel, independent responses to uncoupling.

**Figure 7:**
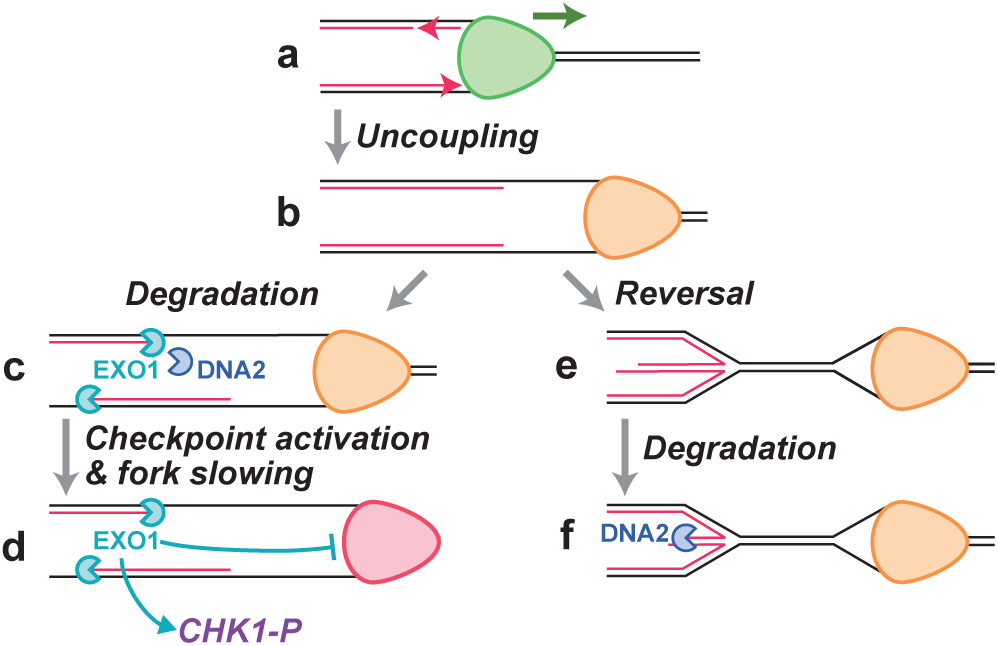
Model for NSD in *Xenopus* egg extracts. **a,** Helicase and polymerase activities are coupled at replication forks. **b,** After DNA polymerases stall, the replicative helicase continues unwinding, producing uncoupling that slows the replisome (orange). **c,** Uncoupling stimulates NSD by EXO1 at uncoupled forks (this study) and DNA2 (shown previously^36^). **d,** Degradation by EXO1 leads to CHK1 phosphorylation and further slows the replisome (red). **e,** Uncoupling triggers fork reversal in parallel to degradation of uncoupled forks. **f,** The regressed arm of a reversed fork undergoes NSD by DNA2 (shown previously ^36^) and not EXO1 (this study).

### 5’-3’ strand-independent degradation of both strands

Our data reveal how a single 5’-3’ degradation process can degrade both strands. The lagging strand is processed from its native 5’ end, and the leading strand from a distinct, distal 5’ entry point (Fig. 3-4). Similar mechanisms may operate in cells, which would explain the predominantly 5’-3’ nuclease activities implicated in degradation^30,31,34,35,39–41^. It is unlikely that the sister fork would act as a 5’ entry point in cells because the sister forks are further apart than in our plasmid system. However, a similar 5’ entry point could be generated through repriming by PrimPol, consistent with the loss of nascent strand degradation upon PrimPol depletion^62^. Additionally, nicks can initiate extensive resection^29,73^ and may serve as an alternative 5’ entry point, which could explain the role of mismatch repair, which culminates in nascent strand nicking, in nascent strand degradation^73^.

Although degradation is strand-independent, the reliance of both strands on a shared degradation factor, EXO1, means that blocking degradation of one strand blocks degradation of the other. Similar reliance on a single degradation mechanism could explain the apparent interdependence of degradation in cells. However, we note that the degradation we studied was at uncoupled, but not reversed, forks and the situation may be different in cells where most degradation depends on fork reversal^1–5^. Indeed, we observed evidence for a residual EXO1-independent degradation mechanism that occurred to a similar extent at all loci (Fig. 4A-D) and was thus distinct, possibly reflecting reversed forks.

### Stability of 3’ ends

Our examination of uncoupled forks revealed stable 3’ ends that did not undergo detectable degradation and instead were slowly extended, even in the presence of multiple polymerase inhibitors (Fig. 3G-H). The simplest explanation is that RFC-PCNA protects the 3’ end, as reported^44,45^. We found that 3’ extension is strongly inhibited by aphidicolin (Fig. S3L-N) and adding ara-dCTP only modestly increases stalling (Fig. 3G-H) suggesting any polymerases at the 3’ end are already inhibited by aphidicolin. Because aphidicolin inhibits B-family DNA polymerases^67^ any polymerase bound at the 3’ end is likely a B-family enzyme such as Pol ε or Pol δ. Whether this polymerase would need to be removed in order to support other repair outcomes, such as template switching^1^, is unclear. However, we note that 3’ end stability and continuous competence for polymerase extension mean that 3’ ends retain the ability to restart for a prolonged period, which would facilitate replication restart following stress.

### Unique role of EXO1 in degradation

It is surprising that degradation of uncoupled forks is predominantly dependent on EXO1, given that DNA2 and MRE11 are both present in our experiments and should also be able to perform degradation. One possibility is that the 5’ RNA at the end of the lagging strand establishes a Ku-dependent barrier to degradation^74^ unless bypassed by EXO1 which can readily degrade a 5’ RNA moiety^75^. However, both MRE11 and DNA2 should be able to cleave beyond the 5’ primer to overcome this barrier^3,75^. It is possible that DNA2 is inhibited by modified nucleotides within the resected strand (e.g. misincorporated ribonucleotides) and that MRE11 is simply not processive enough to result in substantial degradation by itself. Defining which features of uncoupled forks make them EXO1 substrates and whether any additional proteins are required for degradation will require further work.

The Lynch-syndrome associated E109K mutant of EXO1^59^ is MMR proficient^66^, in contrast to other Lynch-syndrome mutations which are thought to cause disease due to MMR defects. Importantly, the E109K mutant is defective in the DNA damage response^66^ but catalytically active^65,76^ raising the possibility of a tumor suppressive role that is distinct from MMR. We found that the *Xenopus* equivalent of E109K abolished nascent strand degradation (Fig. 2E-G). We note that inactivation of MMR proteins can also block nascent strand degradation^73^, which would make nascent strand degradation defects a common link between EXO1 E109K and other Lynch-syndrome mutants. It will therefore be important to further characterize the EXO1 E109K mutant and explore the potential connection of nascent strand degradation to Lynch-syndrome as well as its relationship to EXO1 as a therapeutic target^77^.

### Consequences of nascent strand degradation

The crucial role for EXO1 in ATR activation suggests that helicase-polymerase uncoupling is not sufficient to activate the ATR checkpoint (Fig. 5). It does not reflect a general requirement for EXO1 in ATR activation because ATR-activating DNA structures induced CHK1 phosphorylation independent of EXO1 (Fig. S5D-H). While the mechanistic basis is unclear, one possibility is that degradation of the 5’ end is required to remove a barrier to 9-1-1 clamp loading^54,78^. A role for EXO1 in promoting ATR activation was also reported at post-replicative gaps generated by PrimPol^30^, but the mechanism we observe is distinct because it arises at uncoupled forks and initiates at the 5’ ends of lagging strands rather than internally. Regardless of mechanism, our data show that uncoupling alone is not sufficient for robust ATR activation and that subsequent processing by EXO1 is a crucial step.

That EXO1-mediated degradation actively slows fork progression (Fig. 6) is unexpected. The effect on fork progression is direct because we monitored complete unwinding of the parental template rather than extension of nascent strands (Fig. 1, Fig. 6D-F). It is unclear how lagging-strand degradation at an uncoupled fork would slow progression, especially because impaired lagging-strand synthesis does not slow forks^79,80^. Regardless of the mechanism, we note that EXO1-mediated replication slowing would limit detrimental outcomes of continued fork progression in response to damage while complementing the DNA damage response mounted by EXO1-mediated ATR activation. Thus, the downstream consequences of EXO1 activation that we have identified are well suited to promote genome stability in response to replication obstacles.

## Supplemental figure legends

**Supplemental Figure 1: EXO1 does not impact replication fork uncoupling a.** EXO1-immunodepleted extracts were analyzed by western blotting to determine the extent of EXO1 immunodepletion. **b.** Purified recombinant xEXO1-Flag was analyzed on an SDS-PAGE gel and visualized by Coomassie staining. **c.** Plasmid DNA harboring a LacR array was replicated in Mock- and EXO1-immunodepleted *Xenopus* egg extracts in the absence or presence of purified recombinant xEXO1 protein, and [α-^32^P]dCTP was added to label newly synthesized DNA strands. Once forks were localized to the LacR barrier, aphidicolin and IPTG were added simultaneously to induce uncoupling. **d.** Samples from (c) were separated on an agarose gel and visualized by autoradiography. **e.** Quantification of theta signal from (d). Mean ± s.d. n=3. **f.** The same approach was performed as in (c), but with ara-dCTP instead of aphidicolin to stall nascent strands. **g.** Samples from (f) were separated on an agarose gel and visualized by autoradiography. **h.** Quantification of theta signal from (g). Mean ± s.d. n=3.

**Supplemental Figure 2: EXO1 degradation of uncoupled forks is RAD51-independent a.** Quantification of double-Ys as a % of total signal from Fig. 2b. Mean ± s.d. n=3. **b.** Quantification of reversed forks as a % of total signal from Fig. 2b. Mean ± s.d. n=3. **c.** Quantification of linears as a % of total signal from Fig. 2b. Mean ± s.d. n=3. **d.** Plasmid DNA harboring a LacR array was replicated in Mock- and RAD51-immunodepleted *Xenopus* egg extracts and [α-^32^P]dATP was added to label newly synthesized DNA strands. Once forks were localized to the LacR barrier, aphidicolin and IPTG were added simultaneously to induce uncoupling. Samples were purified and digested as in Fig. 1a. **e.** RAD51-immunodepleted extracts were analyzed by western blotting to determine the extent of RAD51 immunodepletion. **f.** Samples from (d) were separated on an agarose gel and visualized by autoradiography. **g.** Quantification of RI signal from (f). Mean ± s.d. n=4. **h.** Purified recombinant xEXO1 along with E109K and D225A mutants were analyzed on an SDS-PAGE gel and visualized by Coomassie staining. **i.** T=0 samples corresponding to Fig. 2F were separated on an agarose gel and analyzed by autoradiography. **j.** Quantification of Fig. 2F and (i) as in Fig. 1C. Mean ± s.d., n = 3. **k.** Quantification of (i) relative to the buffer condition. Mean ± s.d., n = 3.

**Supplemental Figure 3: 3’ ends undergo slow extension a.** Cartoon depicting the fragment sizes present in the strand-analysis DNA plasmid used to monitor strand-specific degradation using denaturing agarose gels in Fig. 3a-f. Cartoon depicting the control plasmid DNA template, which is linearized by SacI digestion and is analyzed in native agarose gels performed in Fig. 3a-f. **b.** Uncropped native gel of Fig. 3b (top). **c.** Uncropped denaturing gel of Fig. 3b (bottom). **d.** Uncropped native gel of Fig. 3e (top). **e.** Uncropped denaturing gel of Fig. 3e (bottom). **f.** Uncropped denaturing polyacrylamide gel of Fig. 3h (bottom). **g.** Representative histogram of signal intensity in Fig. 3h lane 1, 0 min. **h.** Representative histogram of signal intensity in Fig. 3h lane 2, 30 min. **i.** Representative histogram of signal intensity in Fig. 3h lane 3, 60 min. **j.** Representative histogram of signal intensity in Fig. 3h lane 4, 120 min. **k.** Representative histogram of signal intensity in Fig. 3h lane 5, 240 min. **l.** Plasmid DNA harboring a LacR array was replicated in Mock-immunodepleted *Xenopus* egg extracts. Once forks were localized to the LacR barrier, aphidicolin and IPTG were added simultaneously to induce uncoupling. Samples were purified and digested as in Fig. 3g. **m.** Samples from (l) were separated on a denaturing agarose gel and visualized by autoradiography. **n.** Quantification of the increase in leading strand 3’ end distance over time from (m). Mean ± s.d. n=3. **o.** Plasmid DNA harboring a LacR array was replicated in *Xenopus* egg extracts in the presence of [α-^32^P]dCTP to label newly synthesized DNA strands. Once forks were localized to the LacR barrier, aphidicolin, IPTG, and an excess of unlabeled dCTP nucleotides (“cold chase”) were added simultaneously to induce uncoupling and to analyze potential degradation and resynthesis. **p.** Samples from (o) were purified and digested as in Fig. 3d. Digested samples were separated on native (top) and denaturing (bottom) agarose gels and visualized by autoradiography. **q.** Quantification of leading strand internal fragments from (p). Mean ± s.d. n=3. **r.** Quantification of leading strand end fragments from (p). Mean ± s.d. n=3. **s.** Plasmid DNA harboring a LacR array was replicated in *Xenopus* egg extracts in the presence of [α-^32^P]dCTP and an excess of unlabeled dCTP nucleotides. **t.** Samples from (s) were separated on an agarose gel and visualized by autoradiography. **u.** Quantification of theta signal from (t). Mean ± s.d. n=3.

**Supplemental Figure 4: EXO1 degrades the 5’ ends of both nascent strands a.** Uncropped native gel of Fig. 4b (top). **b.** Uncropped denaturing gel of Fig. 4b (bottom). **c.** Quantification of lagging strand fragments in the mock-immunodepletion condition from Fig. 4b. Mean ± s.d. n=3. **d.** Quantification of lagging strand fragments in the EXO1-immunodepletion condition from Fig. 4b. Mean ± s.d. n=3. **e.** Uncropped native gel of Fig. 4f (top). **f.** Uncropped denaturing gel of Fig. 4f (bottom). **g.** Quantification of leading strand fragments in the mock-immunodepletion condition from Fig. 4f. Mean ± s.d. n=3. **h.** Quantification of leading strand fragments in the EXO1-immunodepletion condition from Fig. 4f. Mean ± s.d. n=3. **i.** Uncropped native gel of Fig. 4k (top). **j.** Uncropped denaturing gel of Fig. 4k (bottom). **k.** Quantification of lagging strand end fragments from Fig. 3b and Fig. 4k. Mean ± s.d. n=3. **l.** Quantification of lagging strand internal fragments from Fig. 3b and Fig. 4k. Mean ± s.d. n=3. **m.** Uncropped native gel of Fig. 4m (top). **n.** Uncropped denaturing gel of Fig. 4m (bottom). **o.** Quantification of leading strand end fragments from Fig. 3e and Fig. 4m. Mean ± s.d. n=3. **p.** Quantification of leading strand internal fragments from Fig. 3e and Fig. 4m. Mean ± s.d. n=3.

**Supplemental Figure 5: EXO1 is specifically required for ATR activation at uncoupled forks a.** Schematic of the assay performed as in Fig. 5a. **b.** pATM from the same samples used in Fig 5C and generated as in (a) was detected by western blotting. **c.** Quantification of pATM signal from (b). Mean ± s.d. n=3. **d.** M13 ssDNA and aphidicolin were added to *Xenopus* egg extracts in the absence or presence of ATRi. **e.** pCHK1 from (d) was detected by western blotting. **f.** Mock- or EXO1-immunodepleted extracts were incubated in the absence or presence of M13 ssDNA. **g.** pCHK1 from (f) were detected by western blotting. **h.** Quantification of pCHK1 signal from (g). Mean ± s.d. n=3.

**Supplemental Figure 6: EXO1 degrades uncoupled forks to restrain fork progression and activate ATR a.** Plasmid DNA harboring a LacR array was replicated in *Xenopus* egg extracts in the presence of Mock or EXO1 antibody and in the absence or presence of purified recombinant xEXO1 protein. [α-^32^P]dCTP was added to label newly synthesized DNA strands. Once forks were localized to the LacR barrier, aphidicolin and IPTG were added simultaneously to induce uncoupling. **b.** Samples from (a) were separated on an agarose gel and visualized by autoradiography. **c.** Quantification of whole-lane signal from (b). Mean ± s.d. n=3. **d.** Quantification of total signal from Fig. 6E. Mean ± s.d. n=3. **e.** Schematic of the assay performed as in Fig. 6D, but in the absence or presence of a cold chase of unlabeled dCTP nucleotides at 40 min. **f.** Samples from (e) were separated on an agarose gel and visualized by autoradiography. **g.** Quantification from (f) of the time in minutes at which fork merger was complete on 50% of molecules for each condition. **h.** Quantification from a biological replicate of (g).

## Methods

### *Xenopus* egg extracts

*Xenopus* egg extracts were prepared from wild-type *Xenopus laevis* male and female frogs (Xenopus1) as previously described^82^. Animal protocols were approved by the Vanderbilt Division of Animal Care and the Institutional Animal Care and Use Committee.

### Plasmid construction and preparation

The construction of pJD145, pJD156, and pJD161 were previously described^36,69^. To create pJD228, a strand-specific digestion scheme was engineered into the pJD161 backbone where digestion with SacI and Nt.BbvCI or Nb.BbvCI would result in 308 nt & 455 nt DNA fragments on either strand. To create pJD229, the same strand-specific digestion scheme was engineered into the pJD161 backbone, along with an additional 2 kb DNA sequence between the 455 nt DNA fragment and the opposing *lacO* array. To create pJD201 and pJD211, the BsrGI/BsiWI DNA fragment from pJD161 (p[lacOx48]) was cloned into pJD228 or pJD229, respectively, that had been digested with BsrGI. To create pJD205, the BsrGI/BsiWI DNA fragment from pJD92 (p[lacOx32]) ^69^ was cloned into pJD228 that had been digested with BsrGI. After DNA replication in extracts, digestion of pJD201, pJD211, or pJD205 with SacI and Nt.BbvCI or Nb.BbvCI results in 308 nt & 455 nt DNA fragments on the nascent leading strand or lagging strand, respectively. After DNA replication in extracts, digestion of pJD201, pJD211, or pJD205 with Nt.BbvCI results in a 308 nt DNA fragment on the nascent leading strand. To create pJD217, all Nt.BbvCI and Nb.BbvCI sites were removed from pJD145.

### Protein purification

Biotinylated LacR protein was expressed and purified as previously described^83^. xEXO1-Flag protein was generated by GenScript. The xEXO1 protein contains a Flag tag on the C-terminus (DYKDDDDK) and was expressed in insect cells and purified by Flag column and Superdex200 column. The xEXO1-Flag storage buffer contains 50 mM Tris-HCl, 500 mM NaCl, 10% glycerol, 1 mM TCEP, pH 8.0.

### DNA replication in *Xenopus* egg extracts

High Speed Supernatant (“HSS”) was prepared by incubating HSS with ATP regenerating system (“ARS”: 20 mM phosphocreatine, 2 mM ATP, and 5 ng/µl creatine phosphokinase) and nocodazole (3 ng/µl) for 5 min at room temperature. To form a replication barrier, 2 volumes of LacR (25.66 µM) were incubated with 1 volume of lacO plasmid DNA (300 ng/µl) for at least 90 minutes at room temperature. To license plasmid DNA, 1 volume of ”licensing mix” was prepared by adding LacR-bound plasmid DNA or plasmid DNA to HSS at a final concentration of 15 ng/µl, followed by incubation at room temperature for 30 min. Nucleoplasmic Extract (NPE) was prepared by supplementation with ARS, dithiothreitol (DTT; final concentration, 2 mM) and [α-^32^P]dATP or [α-^32^P]dCTP. The NPE mix was then diluted to 45% (vol/vol) with 1X egg lysis buffer (ELB; 250 mM sucrose, 2.5 mM MgCl2, 50 mM KCl, 10 mM HEPES pH 7.7). Replication was initiated by adding 2 volumes of NPE mix to 1 volume of licensing mix.

Aphidicolin from *Nigrospora sphaerica* (Sigma Aldrich) was dissolved in DMSO and used at a final concentration of 330 µM (Fig. 5A) except for Fig. 6D-F and Fig. S6E-H where 1 µM was used. Etoposide was dissolved in DMSO and used at a final concentration of 100 µM. Aphidicolin & etoposide were used at a final concentration of 4% (v/v) DMSO in the reaction. ATR inhibitor (ATRi, VE-821) was dissolved in water and used at a final concentration of 10 µM. Mock and EXO1 antibodies were added to reactions at a final concentration of 0.186 mg/ml.

dCTP cold nucleotides were dissolved in water and used at a final concentration of 476 µM. 3X 80-mer DNA oligos containing a 3’ biotin tag were annealed to M13 ssDNA (NEB N4040S) by adding 1 µl of each oligo (500 nM) to 5 µl of M13 ssDNA (250 µg/ml) and incubating at 95°C for 5 min and subsequently allowing to cool to room temperature for 1 h. 80-mer DNA oligo 1 sequence: 5’-ACA AAT AAA TCC TCA TTA AAG CCA GAA TGG AAA GCG CAG TCT CTG AAT TTA CCG TTC CAG TAA GCG TCA TAC ATG GCT TT/Biotin/-3’. 80-mer DNA oligo 2 sequence: 5’-GCT GAT AAA TTA ATG CCG GAG AGG GTA GCT ATT TTT GAG AGA TCT ACA AAG GCT ATC AGG TCA TTG CCT GAG AGT CTG GA/Biotin/-3’. 80-mer DNA oligo 3 sequence: 5’-AGA TTC ACC AGT CAC ACC AGC AGT AAT AAA AGG GAC ATT CTG GCC AAC AGA GAT AGA ACC CTT CTG ACC TGA AAG CGT AA/Biotin/-3’. Annealed M13 ssDNA/3X 80-mer oligos were used in extracts at a final concentration of 2.74 nM.

To induce uncoupling, replication forks were stalled at a LacR-bound lacO array and uncoupled by adding the reaction mix to 10% final reaction volume of IPTG (final concentration, 10 mM in 1X ELB) and Aphidicolin (final concentration, 330 µM in DMSO) or ara-dCTP (final concentration, 450 µM in water). For experiments conducted in Fig. 3H, Aphidicolin & ara-dCTP were both used at a final concentration of 500 µM.

Reactions were sampled into either replication stop solution (8 mM EDTA, 0.13% phosphoric acid, 10% ficoll, 5% SDS, 0.2% bromophenol blue, 80 mM Tris pH 8.0), which stops replication reactions and serves as a DNA loading dye, or extraction stop solution (1% SDS, 25 mM EDTA, 50 mM Tris-HCl pH 7.5), which also stops reactions but allows for DNA purification. All samples were subsequently treated with RNase A (final concentration, 190 ng/µl) and then Proteinase K (final concentration, 909 ng/µl). Samples collected in replication stop were separated by agarose gel electrophoresis at 5 V/cm. Radiolabeled DNA was detected by phosphorimaging and quantification was performed using ImageJ. Samples collected in extraction stop were purified by phenol:chloroform purification and ethanol extraction as previously described^83^.

### Antibodies

Antibody targeting *Xenopus* EXO1 was raised against a polypeptide of LAKACKLLKVANNPDITKVIQKIGQYLKTNITVPEGYIEGFLRANNTFLYQLVFDPVERKLIPLNPYGNDVNPEELNYAGPNMGDSVALQIALGNMDINTRKQIDDYNPDIPQLSHHRSQSWDNKQLNRKTAHTDSIWYTKSEPCKTTKIEEIHSPRGLILPSKK. Antibody targeting *Xenopus* RAD51 was raised against a polypeptide of TTATEFHQRRSEIIQIGTGSKELDKLLQGGIETGSITEMFGEFRTGKTQLCHTLAVTCQLPIDRGGGEGKAMYIDTEGTFRPERLLAVAERYGLSGSDVLDNVAYARAFNTDHQTQLLYQASAMMAESRYALLIVDSATALYRTDYSGRGELSARQMHLARFLRMLLRLADEFGVAVVITNQVVAQVDGAAMFAADPKKPIGGNIIAHASTTRLYLRKGRGETRICKIYDSPCLPEAEAMFAINADGVGDAKD.

### Immunodepletions

Immunodepletions were performed as previously described^84^, but with slight modifications. Protein A-coupled magnetic beads were bound with either 0.5 µg of control, EXO1, or RAD51 antibody per 1 µl of beads. For each round of depletion, 1.29 volumes of antibody-bound beads were incubated with 0.5 volumes of HSS or 1 volume of NPE for 20 min at room temperature with end-over-end rotation. For the EXO1 antibody this was repeated once for HSS and NPE for a total of 2 rounds of depletion. For the RAD51 antibody, this was repeated once for HSS, for a total of 2 rounds of depletion, and twice for NPE, for a total of 3 rounds of depletion. Depleted extracts were collected and used for experiments as described above. For rescue experiments, xEXO1-Flag was added at a final concentration of 188.62 ng/µl.

### NSD assays

To monitor uncoupling and NSD, gel electrophoresis and analysis was performed as previously described ^36,83^. To monitor the replication fork structures formed during NSD, 2D gel electrophoresis and analysis was performed as previously described ^36,83^. To monitor strand-specific degradation, purified NSD intermediates were digested with 0.4 U/µl of SacI and 0.4 U/µl of Nt.BbvCI or Nb.BbvCI in 1X rCutSmart Buffer (NEB) for 1 h at 37°C. Digested products were then separated under native and denaturing agarose gel conditions. For separation under native conditions, 6X native loading buffer (Ficoll, EDTA, SDS, bromophenol blue, and Tris-HCl (1 M, pH 8.0)) was added to samples at a final concentration of 1X before separation on a 0.8% agarose gel at 5 V/cm. For separation under denaturing conditions, digests were stopped by addition of EDTA to a final concentration of 30 mM before addition of 6X alkaline loading buffer (Ficoll, EDTA, xylene cyanol, bromocresol green, and NaOH (10 M)) to a final concentration of 1X. Digests were then separated on a 2% alkaline gel at 1.5 V/cm as previously described ^83^. To monitor leading strand 3’ ends, purified NSD intermediates were digested with 0.4 U/µl of Nt.BbvCI in 1X rCutSmart Buffer (NEB) for 1 h at 37°C. Digested samples were either separated under denaturing agarose gel conditions as described above or were mixed with an equal volume of Gel Loading Buffer II (Invitrogen) to a final concentration of 1X and resolved by a 6% urea-polyacrylamide sequencing gel.

For all NSD assays, radiolabeled DNA was detected by phosphorimaging, and quantification was performed using ImageJ. To measure the stability of leading and lagging strand fragments, denaturing gel fragment signal was normalized to the native gel loading control and expressed as a % of T=0. To measure the extension of the leading strand 3’ end, lane profiles were generated, and each pixel was transformed to scale the signal intensity relative to size. The mode intensity for each timepoint was chosen and size was determined using a standard curve. All 3 repeats were normalized to each other based on the mean of all timepoints from each repeat. The mean of all 3 repeats at time 0 min was set to 0 to measure the extension of the leading strand 3’ end.

Fork merger assay

To quantify the time at which 50% of fork merger had taken place in Fig 6F and Fig. S6G-H linear interpolation was used to determine the time at which θ structures had declined to 50% of total lane signal.

### Western blotting

Western blotting in *Xenopus* egg extracts was performed as previously described ^85^, but with slight modifications. Primary antibodies used for western blotting were: EXO1, RAD51, MCM10 that has been previously described ^86^, pCHK1 (Ser345) (Cell Signaling Technology 2341), and pATM (S1981) (Abcam ab36810). Samples from extract reactions were sampled into Sample buffer. Sample buffer was made by combining 2X SDS buffer (6% SDS, 125 mM Tris-HCl (pH 6.8), 20% glycerol, 0.01% bromophenol blue, 100 mM DTT) and 1X SDS buffer (3% SDS, 62.5 mM Tris-HCl (pH 6.8), 10% glycerol, 0.005% bromophenol blue, 50 mM DTT) in a 1:8 ratio. Samples were separated by 4-12% Bis-Tris gels for MCM10, EXO1, RAD51, and pCHK1 or 4-15% Protean TGX gels for pATM and transferred to PVDF membranes. For western blotting, MCM10, EXO1, and RAD51 blots were incubated in PBST solutions and pCHK1 and pATM blots were incubated in TBST solutions. Membranes were blocked in a solution of 5% milk PBST or TBST for 30 min and then incubated with primary antibody at a dilution of 1:10,000 for MCM10, EXO1, and RAD51, and 1:5,000 for pCHK1 and pATM in 1X PBST or TBST containing 1% BSA for at least 16 h at 4°C. After washing in 1X PBST or TBST, membranes were incubated with horseradish peroxidase (HRP)-conjugated goat anti-rabbit antibody (Jackson Immunoresearch 111-035-003) at a dilution of 1:30,000 for MCM10, EXO1, RAD51, and pATM or 1:10,000 for pCHK1 in 5% milk and 1X PBST or TBST for 1 h at room temperature. Membranes were washed in 1X PBST or TBST and briefly incubated with chemiluminescent HRP detection substrate and developed using the chemiluminescent function on Amersham Imager 600 (GE). Quantification was performed using ImageJ. All 3 repeats were normalized to each other based on the mean of all timepoints from each repeat.

## Acknowledgements

David Cortez (Vanderbilt University School of Medicine) is acknowledged for stimulating discussions.

## Author contributions

JMD conceived of and supervised the project. EJG performed all work except where noted otherwise. OEO performed the experiments in Fig. 5-6 and related supplemental figures with supervision from EJG. TK made the initial observation that EXO1 was crucial for degradation and performed the experiments in Fig. S2D-G. SCC generated the RAD51 antibody used in Fig. S2D-G. JMD and EJG wrote the manuscript.

