## Supplemental Figures for "Strand-independent degradation of uncoupled forks by EXO1 activates ATR and restrains fork progression"

### Supplemental Figure 1

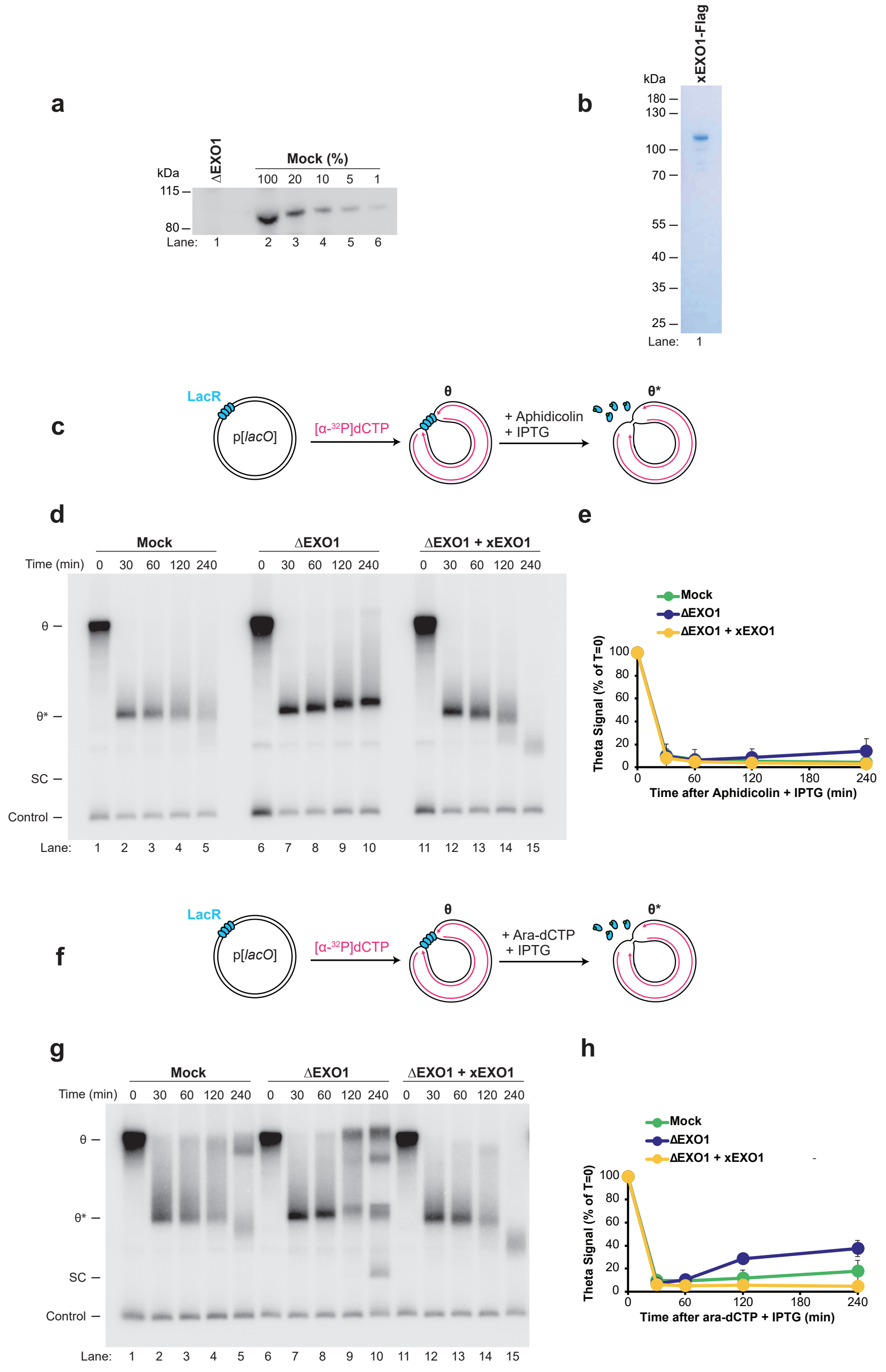

### Supplemental Figure 2

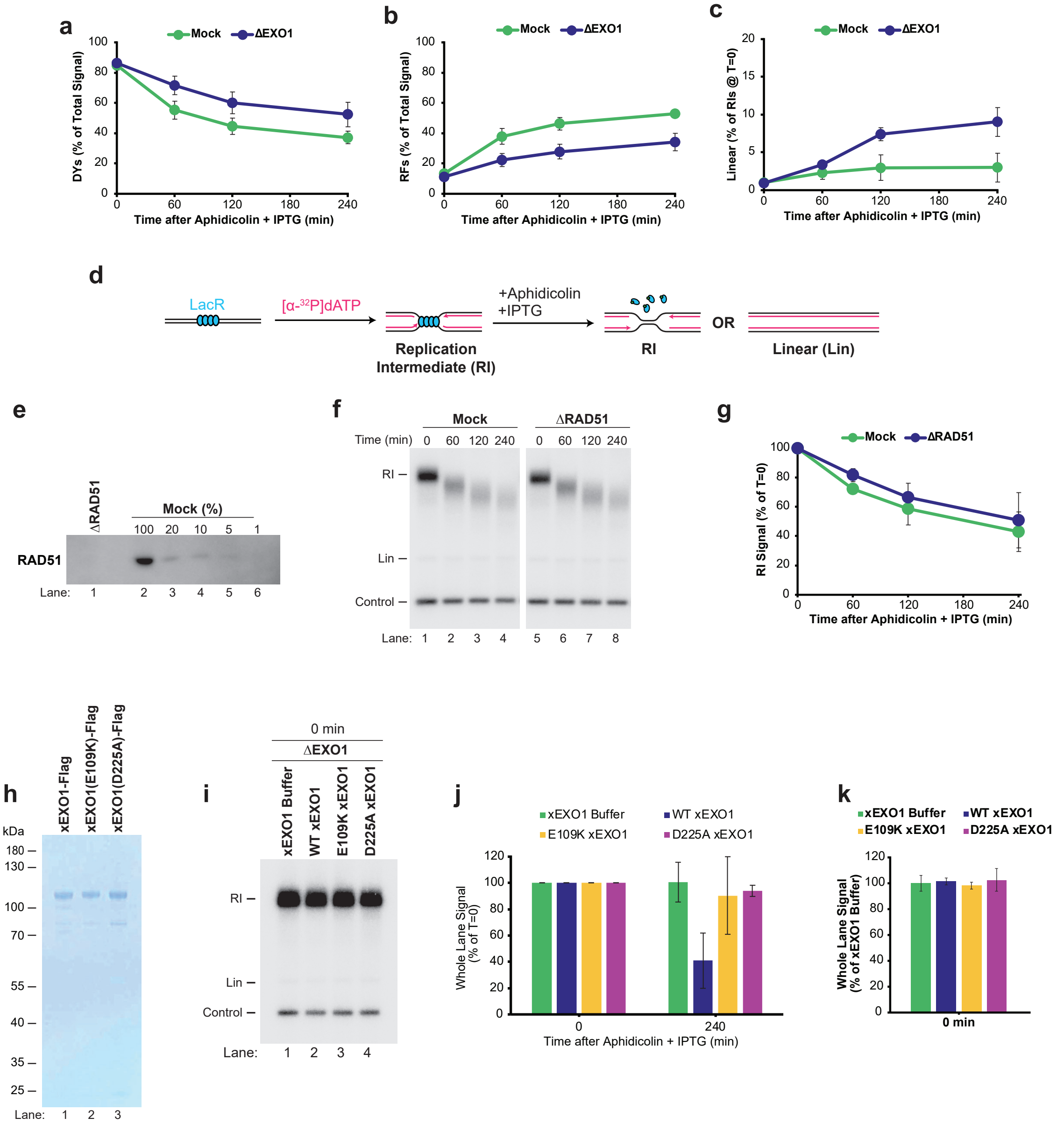

### Supplemental Figure 3

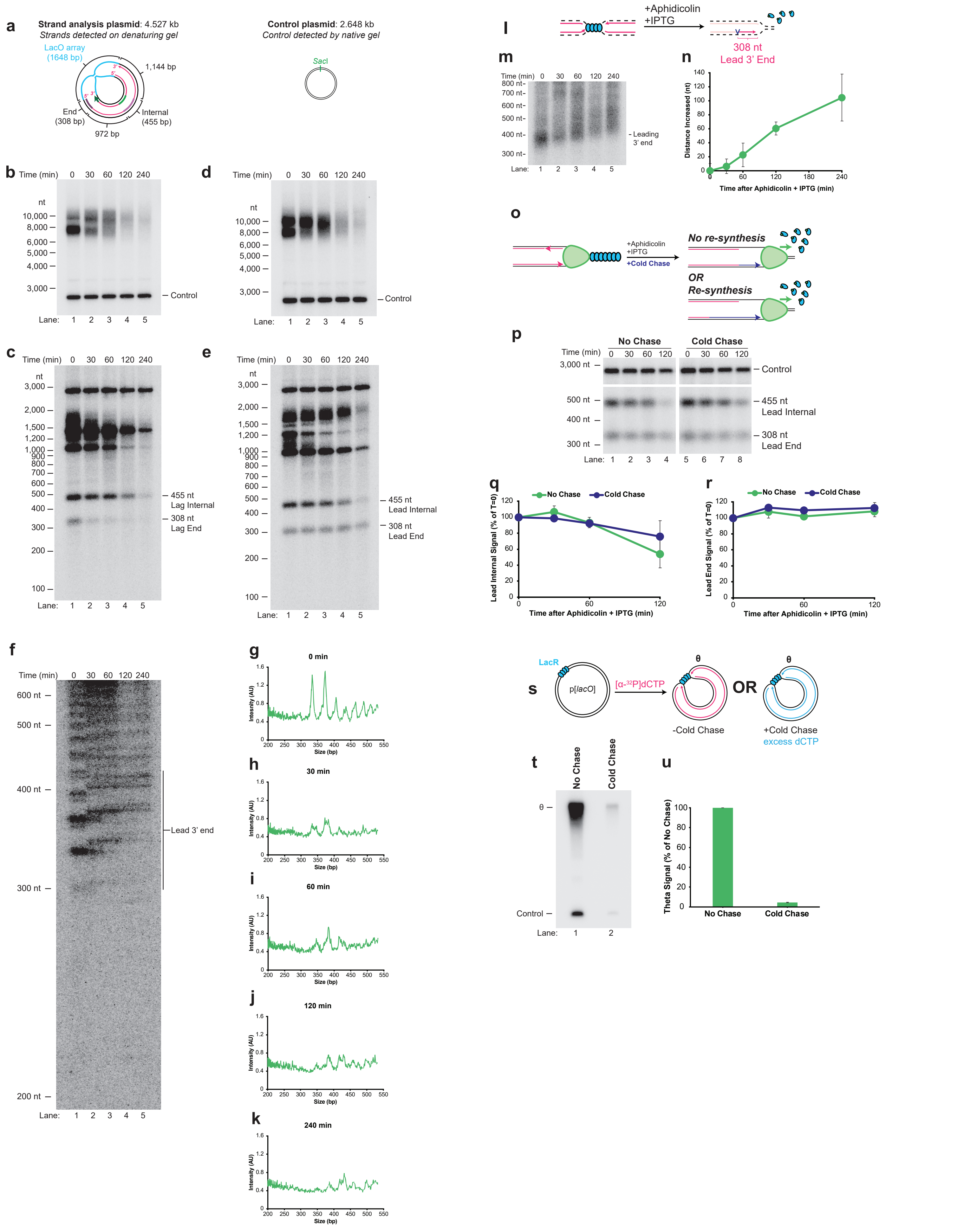

### Supplemental Figure 4

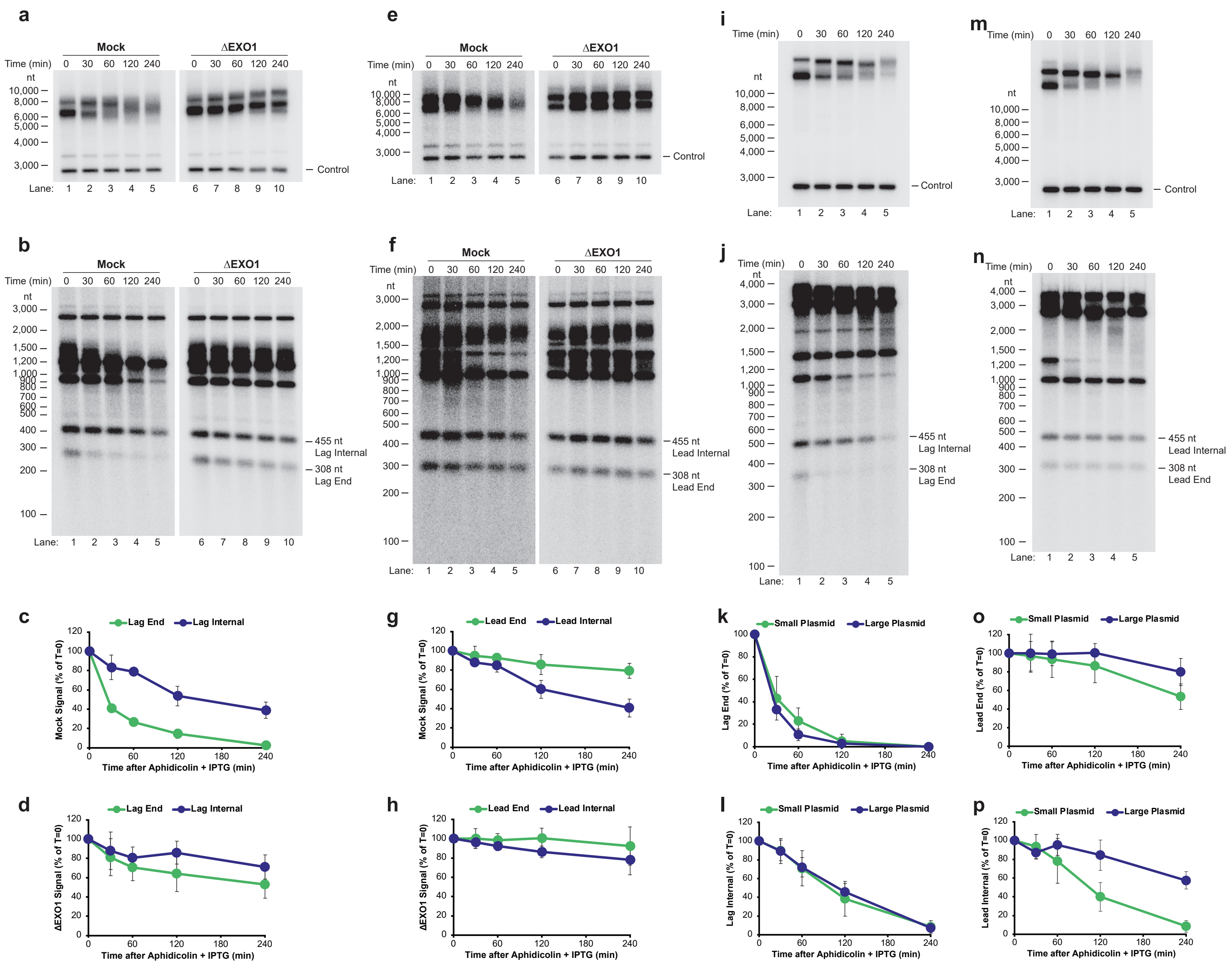

### Supplemental Figure 5

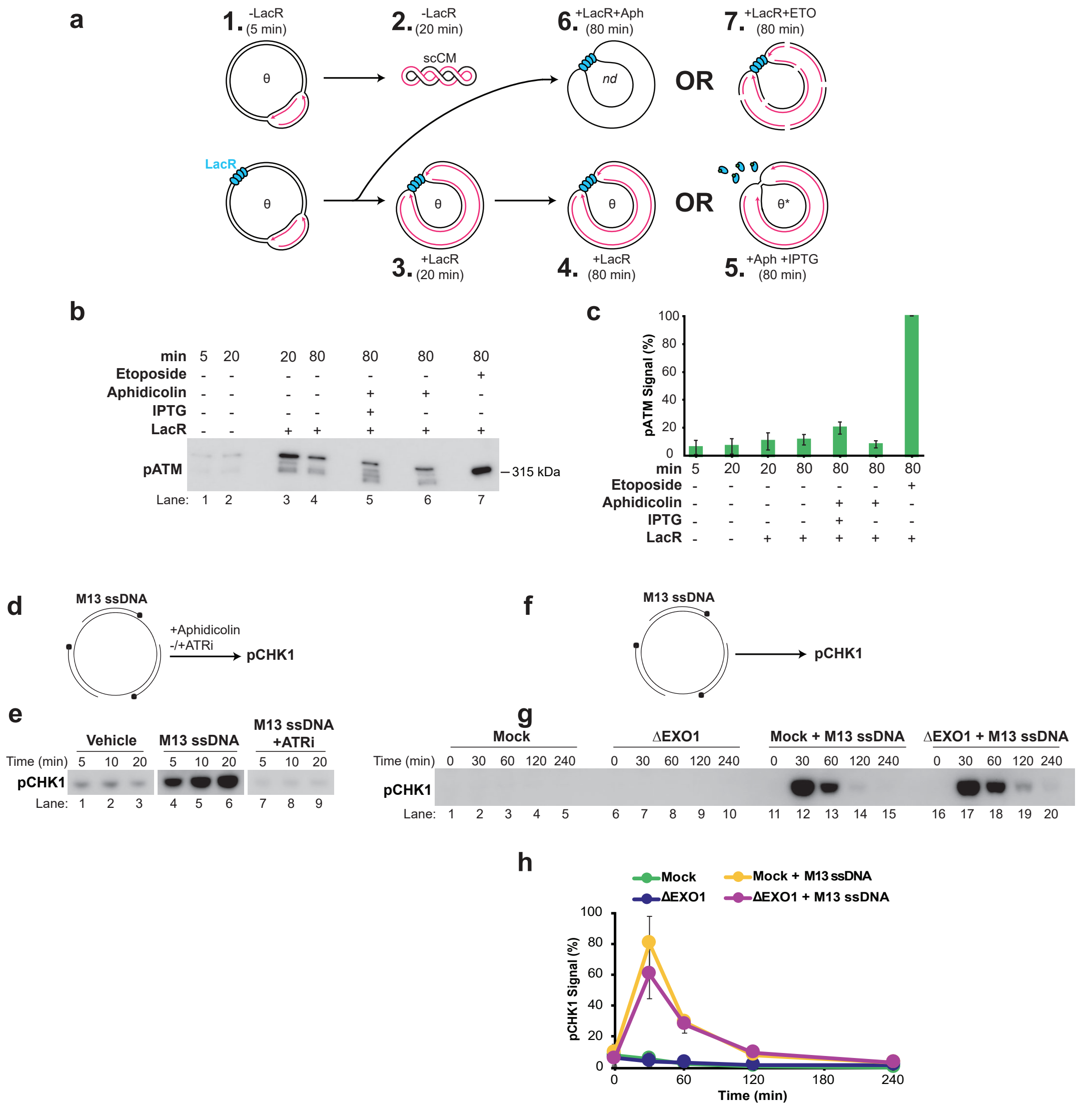

### Supplemental Figure 6

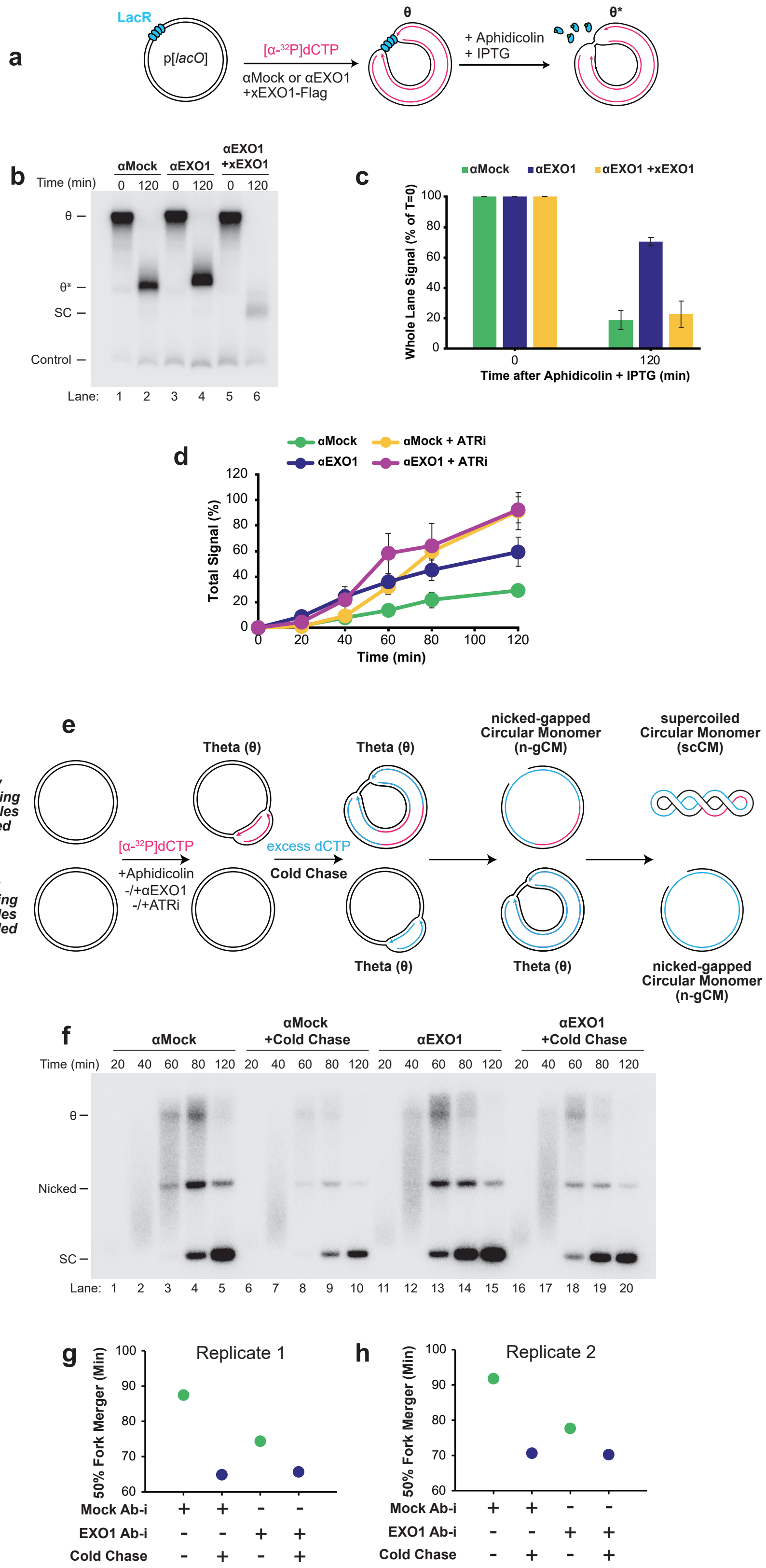
